# W-RESP: Well-Restrained Electrostatic Potential Derived Charges. Revisiting the Charge Derivation Model

**DOI:** 10.1101/2020.09.14.296012

**Authors:** Michal Janeček, Petra Kührová, Vojtěch Mlýnský, Michal Otyepka, Jiří Šponer, Pavel Banáš

**Affiliations:** Department of Physical Chemistry, Faculty of Science, Palacký University, tř. 17 listopadu 12, 771 46, Olomouc, Czech Republic; Regional Centre of Advanced Technologies and Materials, Faculty of Science, Palacký University, tř. 17 listopadu 12, 771 46, Olomouc, Czech Republic; Institute of Biophysics of the Czech Academy of Sciences, Královopolská 135, 612 65 Brno, Czech Republic

## Abstract

Representation of electrostatic interactions by a Coulombic pair-wise potential between atom-centered partial charges is a fundamental and crucial part of empirical force fields used in classical molecular dynamics simulations. The broad success of the AMBER force field family originates mainly from the restrained electrostatic potential (RESP) charge model, which derives partial charges to reproduce the electrostatic field around the molecules. However, description of the electrostatic potential around molecules by standard RESP may be biased for some types of molecules. In this study, we modified the RESP charge derivation model to improve its description of the electrostatic potential around molecules, and thus electrostatic interactions in the force field. In particular, we re-optimized the atomic radii for definition of the grid points around the molecule, redesigned the restraining scheme and included extra point charges. The RESP fitting was significantly improved for aromatic heterocyclic molecules. Thus, the suggested W-RESP(-EP) charge derivation model showed clear potential for improving the performance of the nucleic acid force fields, for which poor description of nonbonded interactions, such as underestimated base pairing, makes it difficult to describe the folding free energy landscape of small oligonucleotides.

## INTRODUCTION

Electrostatics represented by Coulombic interactions of atom-centered partial charges is a fundamental part of molecular mechanics and empirical force field (*ff*) methods.^1^ Although the concept of partial charges may appear to be physically based, there is no quantum chemical observable that corresponds to the partial charges. Thus, their definition is ambiguous and relies on several approximations and assumptions.

Various schemes for partial charge definition have been suggested in the literature. These partial charge derivation methods may be divided into four specific classes.^2–3^ The first class of methods involves partial charges derived directly from experimental data, e.g., κ refinement method,^4^ or from non-quantum mechanical approaches, e.g., TPACM4 method^5^. The second class includes a broad set of methods based on partitioning of the electron charge density (or wave-function) obtained from quantum-mechanical (QM) calculation into atomic populations. Such methods may involve, e.g., the Mulliken population analysis^6^, Löwdin population analysis^7–8^, Hirshfeld population analysis^9–12^, theory of atoms in molecules^13^, population analysis in terms of “fuzzy” atoms^14^, atomic polar tensor-based population analysis^15^ and natural bond orbital population analysis^16–17^. However, the partitioning of the electron density may yield unambiguous and physically ill-defined representation of the molecular dipole or higher-order multiple moments in complex molecules. In addition, the charges obtained by these methods may heavily depend on the QM level used, in particular the choice of basis set. These limitations are addressed in the third class of methods, which are also based on population analyses (mostly the Löwdin population analysis^7–8^), but the charges are subsequently remapped onto a new charge set to reproduce accurately charge-dependent observables, such as the dipole moment, using experimental results or high-level QM calculations.^2^ This class involves, e.g., the charge model (CM) 1, CM2, CM3, CM4, CM4M or CM5.^18–25^ Finally, the fourth class of charge derivation methods is based on reproducing a physical observable predicted from the wave function via fitting of predicted interaction energies^26–28^, dipole moments^29^ or electrostatic potential (ESP)^30–34^.

The charge derivation method based on fitting the ESP is a fundamental methodological feature of the AMBER *ff* families^35^. In contrast to partial charges, the value of the ESP at a particular point in space around a molecule is a QM observable, and thus is physically well defined. In addition, the choice of partial charges reproducing the electrostatic field around a molecule is assumed to also well describe the electrostatic interactions by a Coulomb term involving interactions of the partial charges with electrostatic field of other molecules. The use of ESP-based charges is most likely responsible for the broad success of the AMBER *ff* families in biomolecular simulations. For example, atom-centered ESP charges are amazingly successful in describing base stacking,^36^ although the performance of constant point charges fitted to the ESP is naturally less robust for molecules that sample diverse conformations of the dihedral space^37^

ESP partial charges *q_j_* are obtained by fitting the classical ESP 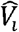 generated by charges *q_j_* to the pre-calculated quantum mechanical molecular ESP *V_i_* evaluated at points *i* around the molecule. For this purpose, the fitting scoring function 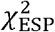 is defined as follows:

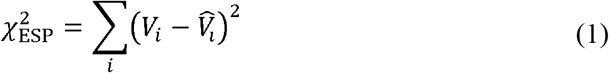

where

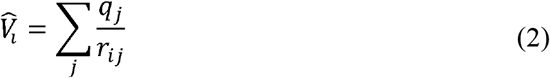

However, despite their success, ESP-derived charges may occasionally be associated with some deficiencies, even disregarding the problems of conformational dependencies of the partial charges. The main problem of ESP fitting is that some charges, especially those corresponding to buried atoms, are statistically ill defined. Beside poor description of buried atoms in larger molecules, the ESP charges may reveal spurious, unphysical conformational dependence and limited transferability between common functional groups in related molecules.^38–42^ These problems are significantly reduced by introducing a restraining term in the fitting scoring function to prevent physically unreasonable values of the fitted charges. Bayly et al.^39–40^ introduced a restrained electrostatic potential (RESP) fitting method and suggested a hyperbolic penalty function 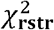 for restraining the charges toward zero value.

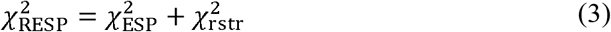

where

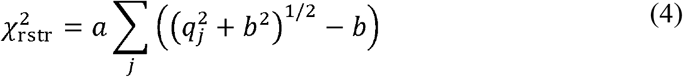

*b* determines the tightness of the hyperbola around its minimum and *a* is the weight factor for the strength of the restraint function.

It was shown that RESP charges lead to smaller variation of the charges among related functional groups.^39–40^ The RESP charges represent the standard charges used in all contemporary AMBER *ff*s. The main purpose of the hyperbolic restraint is to lower the magnitude of the ESP derived charges with only a minimal decrease in the quality of the fitting. Thus, the statistically well-defined charges are only negligibly affected by the restraint, whereas the artificial, statistically ill-defined charges are restrained toward zero value, which precludes or reduces the unintended side effects.

The RESP partial charges perform well for the quantitative aspects of intra- and intermolecular interactions, while having low sensitivity to molecular conformation and configuration.

Recently, we identified deficiencies of nucleic acid force fields which hamper correct description of the structural dynamics and folding of small oligonucleotides.^43^ We showed that this artificial force field behavior is related to inaccurate description of the nonbonded interactions, in particular underestimated base pairing and overestimated sugar-phosphate interactions.^43^ Several groups have attempted to improve the performance of nucleic acid force fields by modification of the nonbonded terms, including modification of van der Waals parameters,^44–48^ combination of van der Waals parameters and partial charges,^49^ or introduction of an additional nonbonded term.^43, 50^ However, as we have argued elsewhere, the potential for improvement of the description of nonbonded interactions by tuning the van der Waals parameters (including NBfix) is rather limited as van der Waals parameters affect the total interaction energy mostly indirectly via electrostatics.^50^ Therefore, despite the abovementioned success of the RESP charge derivation model in AMBER *ff*s, further improvement of the electrostatics is required.

In this study, we revisited the RESP charge derivation method by developing a new approach, denoted as W-RESP, that includes redefinition of the atomic radii for grid construction within the ESP fit and redesigning the method for charge restraint dealing with statistical limitations of the ESP fit. We calibrated the W-RESP method on a set of 47 small bioorganic compounds representing key moieties of biomolecules, i.e., amino acids, nucleotides, lipids and hydrocarbons. We also explicitly probed the effect of extra point (EP) charges on the quality of fit. Finally, the performance of the W-RESP charges was tested by i) probing the effect of the W-RESP charges on the base pairing energy against QM benchmark data, and ii) studying the overall effect of the W-RESP charges on the conformational dynamics of A-RNA duplexes in classical explicit solvent molecular dynamics (MD) simulations.

## METHODS

### Computational details

#### QM calculations and atomic charges

All QM calculations used to derive ESP, RESP and our W-RESP charges were performed using the Gaussian 09 software package^51^. All structures were optimized at the DFT level of theory using the PBE functional for exchange and correlation^52–53^ and the cc-pVTZ basis set^54–55^ with a conductor-like polarizable continuum model of water^56–57^. The ESP around a molecule was calculated at the same level of QM theory, but the effective relative permittivity (dielectric constant) of the solvent model was set to ε=4, as suggested by Duan et al.^58^ The potential was calculated on grid points positioned around the molecule based on the Merz-Singh-Kollman procedure^32^. In contrast to the original procedure, we used 10 shells of points regularly spaced from 1.4 up to 2.0 times the atomic radii (either the original van der Waals radii or our new radii presented in this study) with a density of 17 points/Å^2^. Subsequently, Resp 2.4 software^59–60^ was used to calculate ESP, RESP and W-RESP charges (both with and without EP charges located out of atom centers). For the W-RESP calculation, we implemented a new restraint function in the Resp 2.4 software.

CM5 charges were also calculated using Gaussian 09 based on the same optimized geometries and at the same level of DFT theory with the same implicit solvent settings as the ESP. As there was no straightforward way to calculate the CM5 charges for the EPs (i.e., dummy atoms), we manually redistributed the CM5 charge from a particular heavy atom to its EPs so that the dipole moment of the atom-EPs was the same as for the ESP derived charges, while the total charge of the atom plus its EPs remained the same as the original CM5 charge of the atom alone.

RESP2 charges developed recently by Schauperl et al.^61^ were calculated as described in the original paper using Gaussian 16^62^ and Resp 2.4 software^59–60^. Parameters for the cc-pV(D+d)Z and aug-cc-pV(D+d)Z basis sets^63^ were adopted from https://www.basissetexchange.org/. The density of grid points in each layer was set to 3.0 points/Å^2^.

The quality of the charges was assessed from the standard error (SE) of the ESP fit defined as follows:

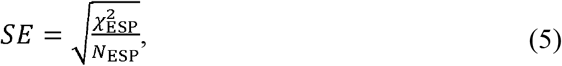

Where 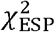 is the scoring function defined in equation (1) and *N_esp_* is number of grid points.

#### Base pairing interaction energies

The performance of the revised W-RESP and W-RESP-EP charges, i.e., set of W-RESP charges with EPs on carbonyls and aromatic imino nitrogens of nucleobases, was initially tested on gas phase interaction energies of base pairing interactions. Molecular-mechanical (MM) interaction energies calculated by different charge sets were benchmarked against the high-level CBS(T) QM method. CBS(T) stands for the MP2^64^/complete basis set limit (CBS) corrected to higher order correlation effects by CCSD(T)^65^ according to a previously described scheme.^66^ We first performed single-point calculations at the MP2/cc-pVTZ and MP2/cc-pVQZ levels to estimate the MP2/CBS energies.^67–68^ Subsequently, the energy difference between the MP2/cc-pVDZ and CCSD(T)/cc-pVDZ calculations was used to estimate a CCSD(T) correction for higher-order correlation effects.^69^ Both the MP2 and CCSD(T) calculations were carried out using the Turbomole 6.3^70–71^ package (http://www.turbomole.com). All calculations using finite basis sets were corrected for the basis set superposition error (BSSE) by a counterpoise correction method.^72^ The final CBS(T) interaction energies were obtained for six conformers of AU, AT and GC base pairs, with the base pair stretch parameter ranging from –0.2 to 0.3 Å. The base pair structures were prepared using our in-house software described in Ref. ^73^ using MP2/cc-pVTZ optimized geometries of the base pair reference frames.

#### MD simulations

The starting topologies and coordinates for classical MD simulations were prepared by using the tLEaP module of the AMBER 16 program package.^74^ The performance of the W-RESP and W-RESP-EP charges was tested on two common RNA duplexes: the r(GCACCGUUGG)_2_ decamer (10-mer), which was excised from the PDB ID 1QC0^75^ structure, and r(UUAUAUAUAUAUAA)_2_ tetradecamer (14-mer, PDB ID 1RNA^76^).

The W-RESP/W-RESP-EP charges of the nucleobase atoms were calculated on N1-methylated pyrimidines and N9-methylated purines. These charges were subsequently transferred into the library files of the standard *ff*99bsc0χ_OL3_^35, 77–79^ RNA *ff* and the charge on the C1’ atom was adjusted to set the total charge of the nucleotide. We preferred to only modify the nucleobase atoms because entire reparametrization of the partial charges in RNA *ff* would have required extensive modification of all the torsion potentials along the RNA backbone. Thus, the only torsion terms affected by the nucleobase atoms’ charge reparametrization and requiring further reparametrization were the glycosidic torsions. In order to adjust the glycosidic torsion potential due to the modified nucleobase charges, we performed MM scans of the glycosidic bond rotation in nucleosides using both standard RESP and modified W-RESP (or W-RESP-EP) charges. Subsequently, we fitted the glycosidic torsion parameters to compensate the effect of W-RESP (or W-RESP-EP) charges on the energy profile of this torsion (see Supplementary Information). Additionally, we used the vdW modification of phosphate oxygens developed by Case et al.^46^, with the affected dihedrals adjusted as described elsewhere.^45^

The RNA duplexes were solvated with the OPC^80^ water model. The minimum distance between the box walls and solute was 12 Å and all simulations were performed in ~0.15 M KCl salt using Joung−Cheatham^81^ ionic parameters. The RNA molecule remained constrained during minimization and optimization of waters and ions. Subsequently, all RNA atoms were frozen and the solvent molecules with counter-ions were allowed to move during a 500-ps long MD run under NpT conditions (*p* = 1 atm., *T* = 298.16 K) to relax the total density. Afterwards, the RNA molecule was relaxed by several minimization runs, with decreasing force constant applied to the sugar-phosphate backbone atoms. Subsequently, the system was heated in two steps: the first step involved heating under NVT conditions for 100 ps, whereas the second step involved density equilibration under NpT conditions for an additional 100 ps. The particle mesh Ewald (PME) method for treating electrostatic interactions was used. Standard unbiased MD simulations were performed under periodic boundary conditions in the NpT ensemble at 298.16 K using a weak-coupling Berendsen thermostat^82^ with coupling time of 1 ps. The SHAKE algorithm, with a tolerance of 10^−5^ Å, was used to fix the positions of all hydrogen atoms, and a 10.0 Å cut-off was applied to non-bonding interactions to allow a 2-fs integration step. The length of MD simulations was 1 μs.

### Structure preparation

Most structures in the training set were drawn from scratch in GaussView 5.0 software^83^. Only the structures of ribose and 2-deoxyribose required specific conformations determined by structural context, and thus were taken from crystal structures of A-RNA, A-DNA and B-DNA (PDB IDs: 464D, 440D and 1BNA, respectively).

Hydroxy, thiol and amino groups bound on an aliphatic alkane chain usually adopt two dominant conformations - anti-periplanar and anticlinal - with respect to the alkyl. Thus, in derivatives of ethane and propane, both these conformations were modeled in the training set.

## RESULTS AND DISCUSSION

### Charge fitting procedure

The fitting procedure described here follows the pioneering studies of Momany^30, 33^, and Kollman and co-workers^31–32, 40^. The atom-centered partial charges were fitted to reproduce the QM ESP at grid points spread around the molecule in a series of shells covered by a given density of points. The charges *q_j_* were obtained by a least squares fit minimizing the following χ^2^ function:

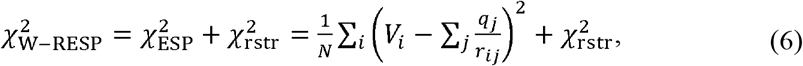

where *N* and *V_i_* correspond to the total number of grid points and QM ESP potential at a particular grid point *i*, respectively, *r_ij_* stands for the distance between grid point *i* and charge fitting center *j*, and 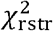 denotes the restraining function.

In contrast to the standard RESP procedure,^40^ the 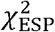 function was normalized by the total number of grid points *N* so that the relative weight between 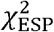 and 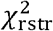 did not depend on the grid density, and thus total number of grid points. Note that increasing the density of the grid in the standard RESP procedure without simultaneously adjusting the weight constant in the restraint term effectively weakens the effect of the restraint, and thus shifts the fit toward pure ESP.

In addition, we employed a different restraint function 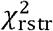 to that used in the standard RESP. The standard RESP procedure uses a hyperbolic restraint toward zero to avoid the statistically ill-defined charges of buried atoms and prevent their artificially large values in magnitude. As suggested by Laio et al., the reliability of the ESP fit might be probed by analysis of the Hessian matrix containing the second derivatives of 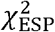 with respect to charges:

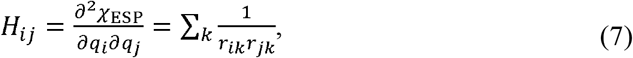

where the summation runs over grid-points *k*. The quality of the ESP fit thus depends only on the geometrical positions of the grid points with respect to the position of charge-fitting centers. Laio et al. showed that whereas the highest eigenvalues of the Hessian matrix corresponding to well-defined narrow minima were associated with lower multipoles (monopole, dipole, quadrupole, etc.), the eigenvectors belonging to the lowest eigenvalues correspond to local changes of few charges without perturbation of low multipoles.^42^ In order to avoid a statistically ill-defined fit due to the broad minima associated with low eigenvalues of the Hessian matrix and simultaneously keep the well-defined parts of the 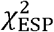 function as unbiased as possible, we suggest using a new weighted harmonic restraint function:

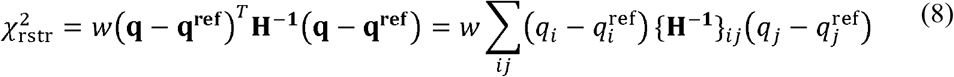

where {*H*^−1^}_*ij*_ are elements of the inverted Hessian matrix **H**, *w* is a scaling coefficient associated with the relative weight of the restraining function 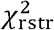 with respect to the 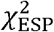 function, and 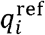 stands for a reference charge set. As the Hessian matrix does not depend on charges, the restraint function 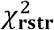 defined above is a parabolic function in the space of fitted charges *q_i_*. It is worth noting that the 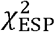 function is parabolic as well and might be rewritten in the following form:

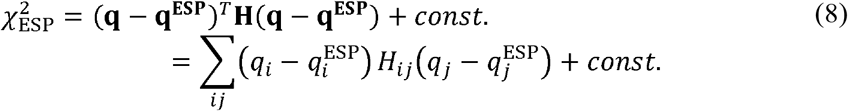

where *H_ij_* and 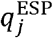 denote elements of the Hessian matrix and fitted ESP charges, respectively. The Hessian matrix **H** and its inverted matrix **H^−1^** share the same eigenvectors and their eigenvalues are inverted with respect to each other. Therefore,whenever the ESP charges are well-defined by large eigenvalues of the Hessian matrix **H** corresponding to high curvature of the 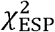 function and narrow minima in the charge space, the restraint function is weak and negligible. Thus, the charges are negligibly biased by the restraint in this case. Conversely, in the case of ill-defined charges due to small eigenvalues of matrix **H** resulting in small curvature and broad minima of the 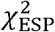 function, the restraint function is strong and shifts the ill-defined charges toward the chosen physically reliable reference charges *q*^ref^. The asymptotic behavior of such a restraint fit with respect to the scaling coefficient *w* makes the fitted charges converge to pure ESP charges or *q*^ref^ charges for *w* approaching zero and infinity, respectively. In the case of finite *w*, the fitted charges are close to ESP values whenever the charge is well-defined by the 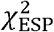 function or are otherwise shifted to alternative physically reliable *q*^ref^ charges. Because of this behavior, the present charges will be referred to as “well-restrained ESP” charges (W-RESP). As pointed out by Laio et al., the parabolic restraint to physically reliable non-zero charges *q*^ref^ avoids the biasing of the atomic partial charges in charged groups such as carboxyl group or phosphate moiety toward unphysical, neutral charges.^42^ An important advantage of the parabolic form of the fitting function is that the optimal charges can be solved analytically by solving a set of linear equations with particular constraints, e.g., for the total charge of the molecule.

In this study, the reference charges *q*^ref^ were represented by CM5 charges^3^ derived from the Hirshfeld population analysis^9^ with improved accuracy for prediction of dipole moments. Thus the CM5 charges provided a physically reliable reference charge set as their values were directly related to the charge density and represented an accurate estimate of low multipole moments.

### Training set of small molecules

In order to parameterize the W-RESP method, in particular to find an optimal value for the weighting parameters *w*, we built up a dataset containing 47 molecules covering all fragments needed for modeling nucleic acids, proteins and small organic compounds. For the nucleic acid fragments, we prepared 14 molecules covering three distinctive chemical sub-units: five-carbon sugar molecules (ribose or deoxyribose), nitrogenous bases, including N-methylated nucleobases, and phosphate groups. The peptide fragments included two subsets covering backbone (N-methylacetamide), and side chains molecules. The final part of the training set comprised four small organic molecules that could not be classified as amino acids or nucleic acid fragments (acetone, dimethylamine, dimethyl ether, and dimethyl sulfide).

**Figure 1.**
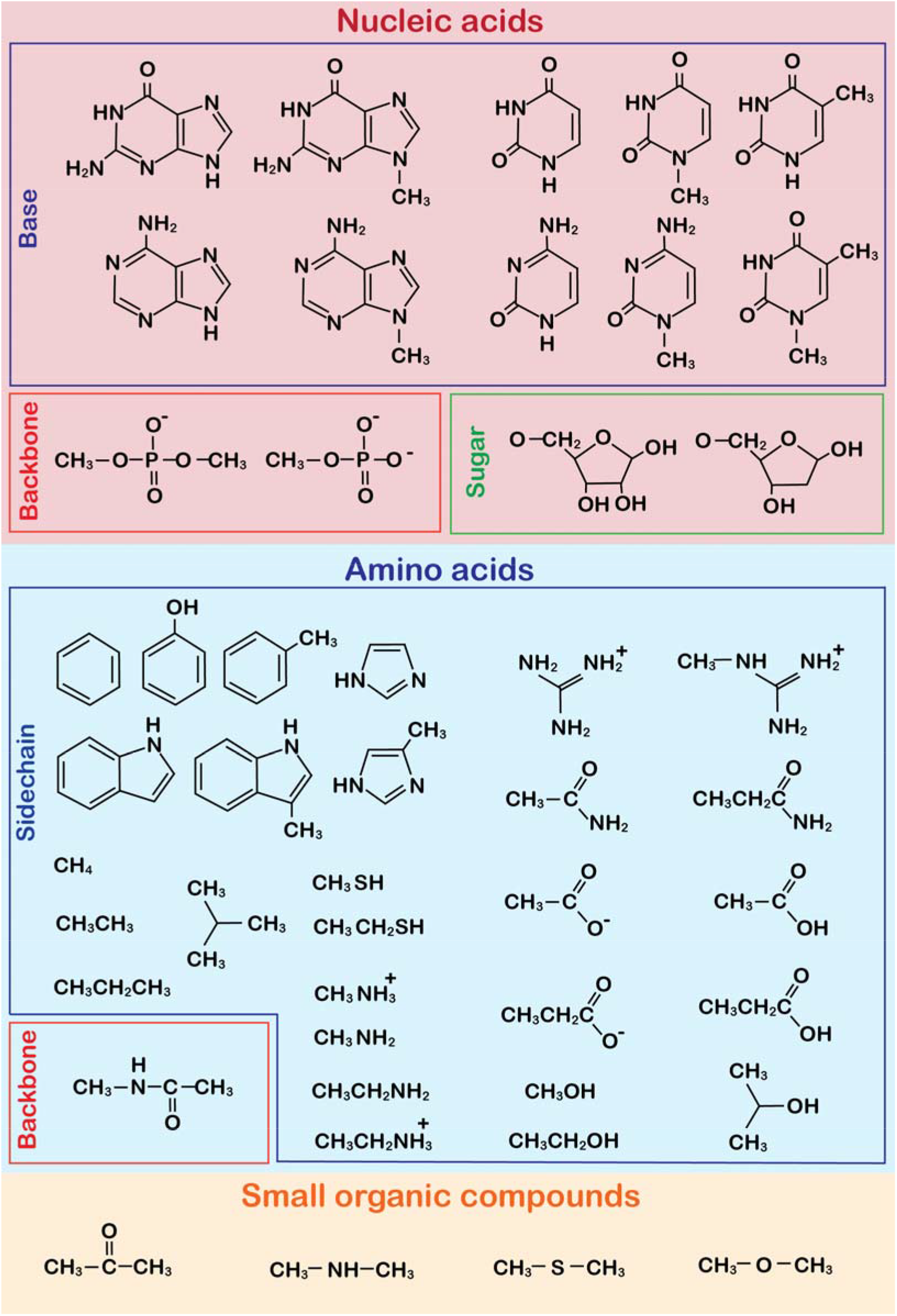
Training set of small molecules used for W-RESP parametrization. The dataset was divided into three main subsets covering nucleic acids, amino acids and other small organic compounds.

### Atomic radii and grid definition

Kollman and co-workers showed that grid points generated on shells with a particular density of points provide sufficient sampling of the ESP potential around molecules providing all points lie outside the van der Waals radii of atoms.^31–32^ We analyzed the grid obtained by the standard Merz-Singh-Kollman procedure^32^ for the set of small molecules used in this study and found that in some cases, mainly for aromatic molecules, some of the grid points were deeply buried in the molecular electron density. Such points showed high residual differences between the QM and MM ESP potential, and thus seemed to represent influential points for the fit. The QM ESP potential of these points was significantly affected by their deep penetration into the electron density. Hence, these points sample regions that cannot be fitted into atom-centered charges and should be considered as outliers.

This effect can be demonstrated by the ESP fit of a benzene molecule. The Merz-Singh-Kollman radius of the carbon atoms is 1.5 Å. Therefore, the innermost shell of grid points lying at 1.4 times the radius is at a distance of 2.1 Å from the carbons. This suggests the grid points reach an area located 1.6 Å above and below the center of the ring plane. Such a region is assumed to be never sampled by any MM atom in MD simulations (note that a typical stacking distance is ~3.2-3.4 Å and even in a T-shape interaction, the hydrogen is not closer to the ring center than ~2.1 Å).

Furthermore, as shown in Figure 2, these points are located in a region where QM ESP gradients cannot be reproduced by atom-centered MM charges. These influential outlier points are located along the normal axis of the benzene ring. Therefore, their ESP potential is mainly affected by the axial part of the molecular quadrupole moment. However, the fit tends to compensate the discrepancy between the MM and QM ESP at these points by tuning the in-plane distribution of MM charges, which is thus biased in an unphysical, artificial way.

**Figure 2.**
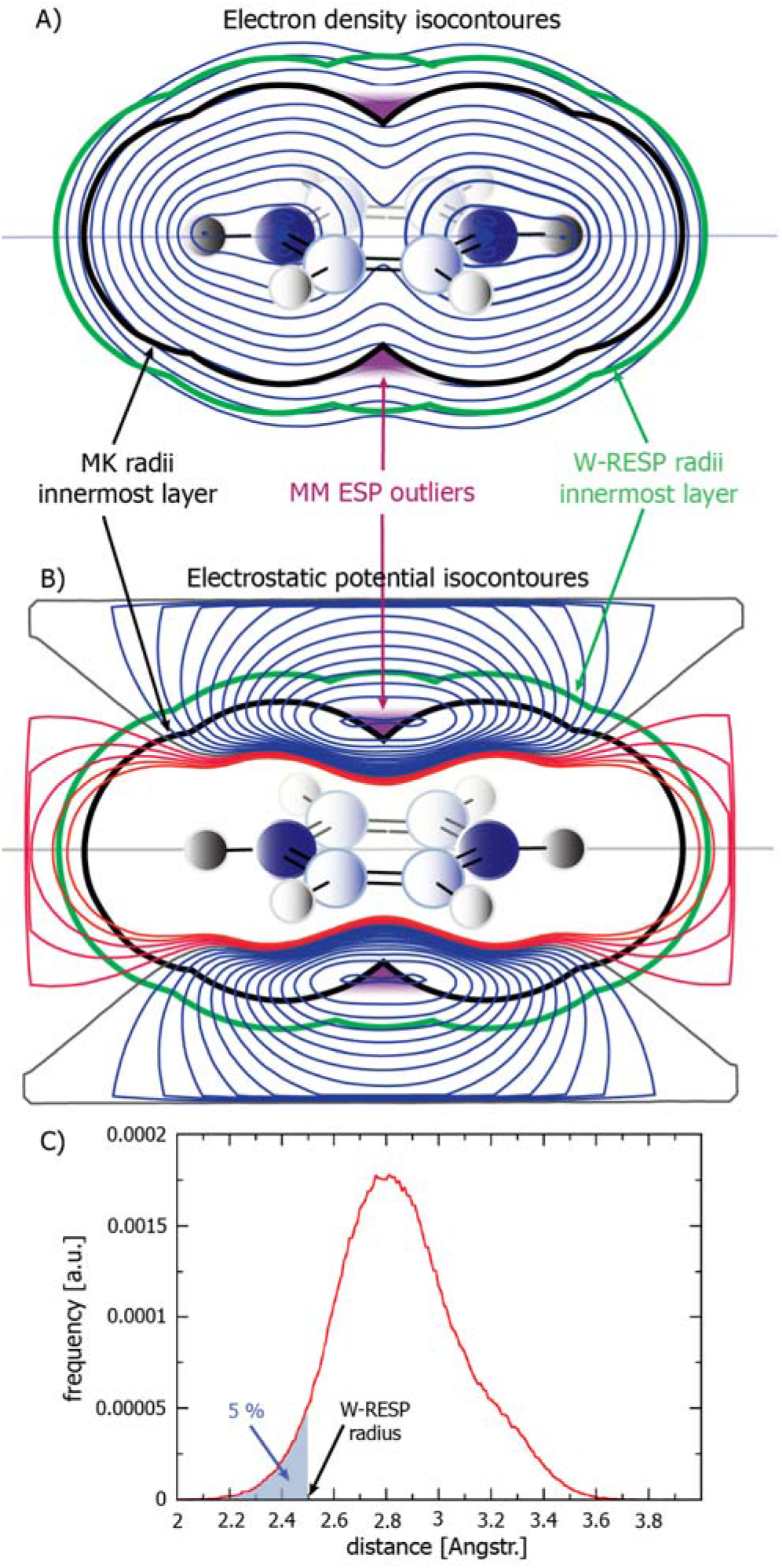
Isocontours of A) the electron density and B) electrostatic potential (ESP) in a plane perpendicular to the benzene ring. The black and green thick contours represent the innermost layers of the grid points around the benzene molecule as generated by standard Merz-Singh-Kollman and re-defined W-RESP radii, respectively. The locations of the influential outlier points buried in the electron density are highlighted in purple. C) Population of the minimal distance of any MM atom from the aromatic carbons in MD simulations of r(gcGAGAgc) and r(gcUUCGgc) RNA oligonucleotides using the AMBER *ff*. The W-RESP radii are defined so that the probability of any MM atom to sample a region inside the innermost grid point layer defined by these radii is as low as 5%, i.e., this region is not sampled by any MM atom with 95% probability.

To eliminate these influential outlier grid points above and below the centers of aromatic rings, we suggest using an additional sphere with radius of 1.89 Å in the center of each 5- or 6-membered ring for defining the grid point layers so that the grid points along the normal axis of the aromatic rings are not closer than 2.65 Å to the center of the ring. Besides exclusion of the grid points buried inside the molecular electron density, the quality of the ESP fit can be further improved by positioning grid points in regions that are often sampled by other MM atoms in the MD simulations so that the electrostatic field generated by MM atoms is sufficiently accurate in these particular regions of interest. To find the optimal positioning of the grid points, we measured the distribution of the minimal distances between atoms of a given atomic type and any MM atoms around them in the set of MD simulations. For this purpose, we reanalyzed simulations of r(gcGAGAgc) and r(gcUUCGgc) RNA tetraloops published in Ref. ^50^. Subsequently, we calculated 5% quantiles for all distributions corresponding to each atomic type, which were used to define the distance of the innermost layer from the atoms (Figure 2c). In other words, the space inside the innermost grid layer was only rarely visited by any MM atom in the MD simulations (with probability 5%), and therefore did not need to be involved in the ESP fit. Note that the standard Merz-Singh-Kollman scheme defines the distance of the innermost grid point layer as 1.4 times the Merz-Kollman radii. Therefore, we calculated new atomic radii from the above mentioned 5% quantile distances by dividing these values by 1.4. Henceforth, we denote these radii as W-RESP radii (see Table 1 for comparison of standard Merz-Kollman and suggested W-RESP radii, respectively). The grid derived by the W-RESP radii and additional sphere in the centers of 5- and 6-membered aromatic rings was labeled as the W-RESP grid.

**Table 1:**
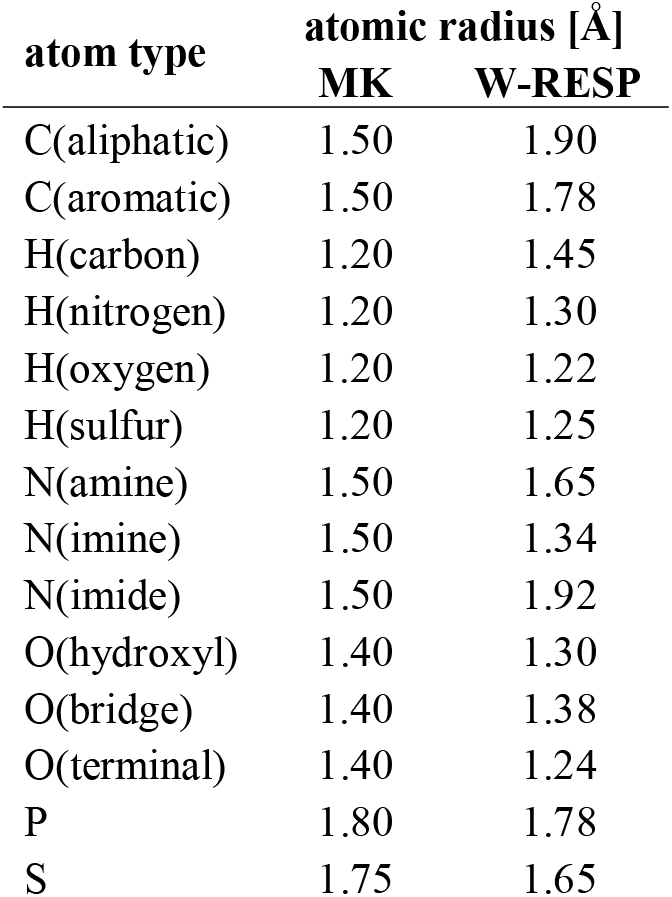
Comparison of the standard Merz-Kollman (MK) and developed W-RESP atomic radii for different atom types.

| atom type | atomic radius [Å] |  |
| --- | --- | --- |
|  | MK | W-RESP |
| C(aliphatic) | 1.50 | 1.90 |
| C(aromatic) | 1.50 | 1.78 |
| H(carbon) | 1.20 | 1.45 |
| H(nitrogen) | 1.20 | 1.30 |
| H(oxygen) | 1.20 | 1.22 |
| H(sulfur) | 1.20 | 1.25 |
| N(amine) | 1.50 | 1.65 |
| N(imine) | 1.50 | 1.34 |
| N(imide) | 1.50 | 1.92 |
| O(hydroxyl) | 1.40 | 1.30 |
| O(bridge) | 1.40 | 1.38 |
| O(terminal) | 1.40 | 1.24 |
| P | 1.80 | 1.78 |
| S | 1.75 | 1.65 |

In order to compare the quality of the ESP fit using grid points defined by the standard MK and developed W-RESP radii, we calculated the SE of the ESP fit (see equation 5) for all 47 molecules in the training set (see Methods section). Both sets of ESP charges were calculated with equivalencing of chemically identical atoms in two stages similarly to the procedure used in the RESP scheme. To enable fair comparison of both charge sets, SEs were calculated on the same W-RESP grid comprising 10 layers of grid points with 17 points/Å^2^. Thus, the MK ESP charges were calculated using the grid defined by MK radii, but the SEs were subsequently recalculated using the W-RESP grid. The quality of the ESP fit was visibly improved by using the W-RESP radii for aromatic compounds, such as benzene, indole and their methyl-derivates, but also for methane. A modest improvement was also observed for nucleobases, deprotonated carboxylic acids, methylated phosphate and compounds containing planar amino group Table 2). For a complete comparison, see Supporting Table S1 in the Supporting Information.

**Table 2:** Standard error of the ESP fit (as defined in equation 5), minimal (min) and maximal (max) difference between the MM and QM ESP potential at the grid points 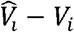 (see equation 1) in ×10^−3^ Hartree/e^−^, all calculated on the same W-RESP grid.

|  | MK radii |  |  | W-RESP radii |  |  | reduction<br>of std error<br>[%] |
| --- | --- | --- | --- | --- | --- | --- | --- |
| | Std<br>error | Min<br>$\hat{V}_i - V_i$ | max<br>$\hat{V}_i - V_i$ | Std<br>error | Min<br>$\hat{V}_i - V_i$ | max<br>$\hat{V}_i - V_i$ | |
| toluene | 0.39 | -2.09 | 2.08 | 0.30 | -2.93 | 1.32 | 22.3 |
| indole | 0.41 | -4.03 | 1.88 | 0.33 | -3.87 | 1.85 | 20.2 |
| methane | 0.17 | -1.02 | 0.58 | 0.14 | -0.95 | 0.43 | 19.2 |
| benzene | 0.50 | -2.18 | 1.33 | 0.40 | -3.54 | 1.12 | 18.9 |
| 3-methylindole | 0.43 | -2.82 | 2.40 | 0.36 | -2.72 | 2.18 | 18.1 |
| methyluracil | 0.62 | -7.24 | 3.98 | 0.54 | -5.20 | 4.73 | 12.0 |
| thymine | 0.68 | -7.60 | 4.59 | 0.60 | -5.48 | 5.08 | 11.8 |
| uracil | 0.75 | -7.56 | 4.37 | 0.67 | -5.52 | 4.96 | 11.7 |
| acetamide | 0.61 | -7.92 | 3.87 | 0.54 | -6.24 | 4.27 | 11.2 |
| methylcytosine | 0.57 | -7.67 | 3.32 | 0.51 | -5.98 | 2.95 | 11.2 |
| cytosine | 0.75 | -9.34 | 5.29 | 0.68 | -7.21 | 6.14 | 9.3 |
| guanidinium | 0.33 | -2.82 | 1.00 | 0.30 | -2.43 | 1.06 | 8.7 |
| methylphosphate | 1.19 | -7.96 | 5.34 | 1.09 | -8.37 | 4.68 | 8.4 |
| methylguanine | 0.82 | -7.70 | 8.10 | 0.75 | -8.66 | 5.72 | 8.1 |
| guanine | 0.84 | -7.69 | 7.72 | 0.78 | -8.44 | 6.20 | 7.3 |
| acetate | 1.18 | -10.07 | 5.37 | 1.10 | -7.90 | 5.42 | 6.7 |
| propionate | 1.19 | -10.14 | 8.73 | 1.12 | -7.71 | 9.07 | 5.8 |
| dimethylphosphate | 1.04 | -7.38 | 5.34 | 0.98 | -7.45 | 4.15 | 5.3 |
| propionic acid | 0.75 | -6.61 | 5.58 | 0.71 | -5.54 | 5.79 | 5.3 |

### Extra point charges

A well-known intrinsic limitation of the atom-centered point charge model is its poor description of the anisotropic charge distribution around atoms carrying lone pairs.^84^ This limitation was more pronounced for charges derived using the W-RESP grid as the W-RESP radii of some polar atoms were even smaller than the corresponding MK radii Table 1).Thus, the ESP fit with W-RESP charges was more sensitive to the description of the anisotropic charge distribution around these polar atoms. This limitation can be partially solved by two different approaches: introduction of additional point charges (extra points, EPs) to the model^85–86^ or by explicitly including multipoles in the force field^87^. To retain the simplicity and efficiency of the classical pair-additive force field, we focused on the former approach. Several previous studies have discussed the ideal number and position of EPs^85–86, 88–90^. Here, we optimized these parameters using the broad set of 47 molecules in combination with W-RESP scheme.

As in similar studies, see e.g. Ref. ^85^, we aimed to find the optimal number and position of EPs by minimizing the SE (equation 5) of the ESP fit while keeping the magnitude and position of the EP charge within a reasonable range suitable for MD simulations. For each functional group occurring in the training set of 47 molecules (see Methods section), one or two EPs were placed at different positions around the polar heavy atoms. The distance between the EPs and heavy atom was scanned over the range 0.2 to 0.5Å (or up 0.7 Å in case of sulfur). The angular degree of freedom represented by the C-X-EP angle and Y-C-X-EP torsional angle was scanned in steps of 5 degrees. We considered only configurations of EPs that did not break the symmetry of the functional group. This generated N-dimensional maps of the relative SE (normalized to the ESP SE of the fit without EPs) as a function of the EP position. These maps of a particular functional group, e.g., imino nitrogen, corresponding to different molecules in the training set of 47 molecules were subsequently averaged to obtain the optimal position of EPs for the particular functional group Figure 3).

**Figure 3:**
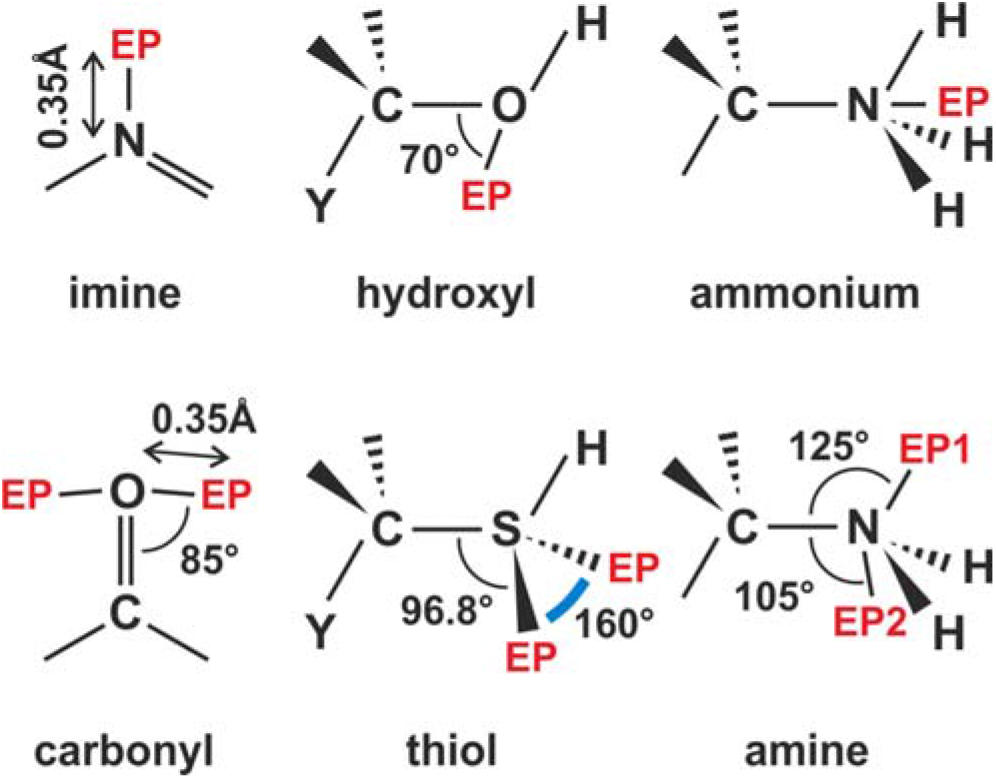
Optimal positions of extra point charges of functional groups contained in the training set. All EPs were located 0.35 Å from the central heavy atom with the exception of the thiol group, where they were 0.7 Å away from the sulfur. For details of the optimization of the EP positions, see the Supporting Information. The positions of EPs on sulfur were adopted from the study of sulfur organic compounds by Yan et al.^90^

The identified optimal positions of EPs Figure 3, Tables S2-S13) agreed well with those suggested in previous studies^85–86, 88–90^ The most significant effect of EPs was observed for heterocyclic aromatic compounds, in particular nucleobases Table 3). Significant improvement was also observed for molecules with amine and thiol groups (Tables S9,S7) and the majority of compounds carrying imine, carbonyl and ammonium groups (Tables S2,S3,S11). On the other hand, the effect of additional EPs was only modest for compounds containing a hydroxyl group (Table S5).

**Table 3:** Effect of extra point charges on the ESP fit (equivalenced in two RESP-like stages) using the W-RESP grid as measured by the SE of the fit, minimal (min) and maximal (max) difference between the MM and QM ESP potential at the grid points 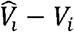 in ×10^−3^ Hartree/e^−^.

|  | No extra points |  |  | Extra points |  |  | reduction<br>of std error<br>[%] |
| --- | --- | --- | --- | --- | --- | --- | --- |
| | Std<br>error | min<br>$\hat{V}_i - V_i$ | max<br>$\hat{V}_i - V_i$ | Std<br>error | min<br>$\hat{V}_i - V_i$ | max<br>$\hat{V}_i - V_i$ | |
| nucleic acids |  |  |  |  |  |  |  |
| adenine | 0.82 | -8.46 | 5.99 | 0.38 | -4.28 | 1.74 | 53.7 |
| guanine | 0.78 | -8.44 | 6.20 | 0.43 | -4.83 | 1.89 | 44.9 |
| uracil | 0.67 | -5.52 | 4.96 | 0.38 | -2.98 | 1.13 | 43.5 |
| thymine | 0.60 | -5.48 | 5.08 | 0.38 | -3.83 | 2.08 | 35.7 |
| cytosine | 0.68 | -7.21 | 6.14 | 0.38 | -3.31 | 1.49 | 44.9 |
| methyladenine | 0.79 | -8.52 | 6.50 | 0.37 | -4.64 | 1.54 | 52.8 |
| methylguanine | 0.75 | -8.66 | 5.72 | 0.42 | -4.73 | 1.80 | 43.9 |
| methyluracil | 0.54 | -5.20 | 4.73 | 0.28 | -2.53 | 1.11 | 48.1 |
| methylthymine | 0.56 | -4.82 | 4.89 | 0.31 | -2.55 | 2.78 | 44.0 |
| methylcytosine | 0.51 | -5.98 | 2.95 | 0.34 | -3.16 | 1.42 | 32.7 |
| amino acids |  |  |  |  |  |  |  |
| imidazole | 1.16 | -6.69 | 10.05 | 0.49 | -4.26 | 1.89 | 57.8 |
| 4-methylimidazole | 0.97 | -6.69 | 7.33 | 0.42 | -4.11 | 1.99 | 57.2 |
| acetic acid | 0.81 | -5.33 | 6.57 | 0.49 | -5.17 | 1.89 | 38.9 |
| acetamide | 0.54 | -6.24 | 4.27 | 0.44 | -3.22 | 1.46 | 18.5 |
| propionic acid | 0.71 | -5.54 | 5.79 | 0.50 | -3.98 | 3.59 | 29.6 |
| methylammonium | 0.85 | -3.24 | 4.69 | 0.29 | -2.34 | 1.69 | 65.5 |
| small organic molecules |  |  |  |  |  |  |  |
| acetone | 0.68 | -7.12 | 6.21 | 0.26 | -3.15 | 0.91 | 62.3 |

### Optimization of the restraint’s relative weight in the W-RESP scheme

The scaling coefficient *w* corresponding to the relative weight of the restraint term was scanned over the range 10^−15^ to 1. All W-RESP fitting was performed with equivalencing of the charges of chemically identical atoms in a two stage model as suggested by Bayly et al. ^39–40^. The optimal value of the scaling coefficient *w* was evaluated using the training set of 47 molecules Figures 1, 4 and S12,S13) either with or without EPs, henceforth labeled as W-RESP and W-RESP-EP, respectively. Similarly, as in the original RESP study by Bayly et al. ^39–40^, the optimal value of the coefficient *w* was set to restrain the charges of buried atoms, while the overall quality of the fit was not significantly affected by the restraint, i.e., the SE of the W-RESP fit was still comparable to its ESP value. Based on these criteria, a value of 10^−13^ was found to be optimal for *w*. Higher values resulted in a significantly lower quality of fit Figures 4 and S12, S13). The optimal value of the coefficient *w* provided a significant restraining effect on statistically ill-defined charges of the buried atoms, as shown for the acetone molecule Table 4).

**Figure 4:**
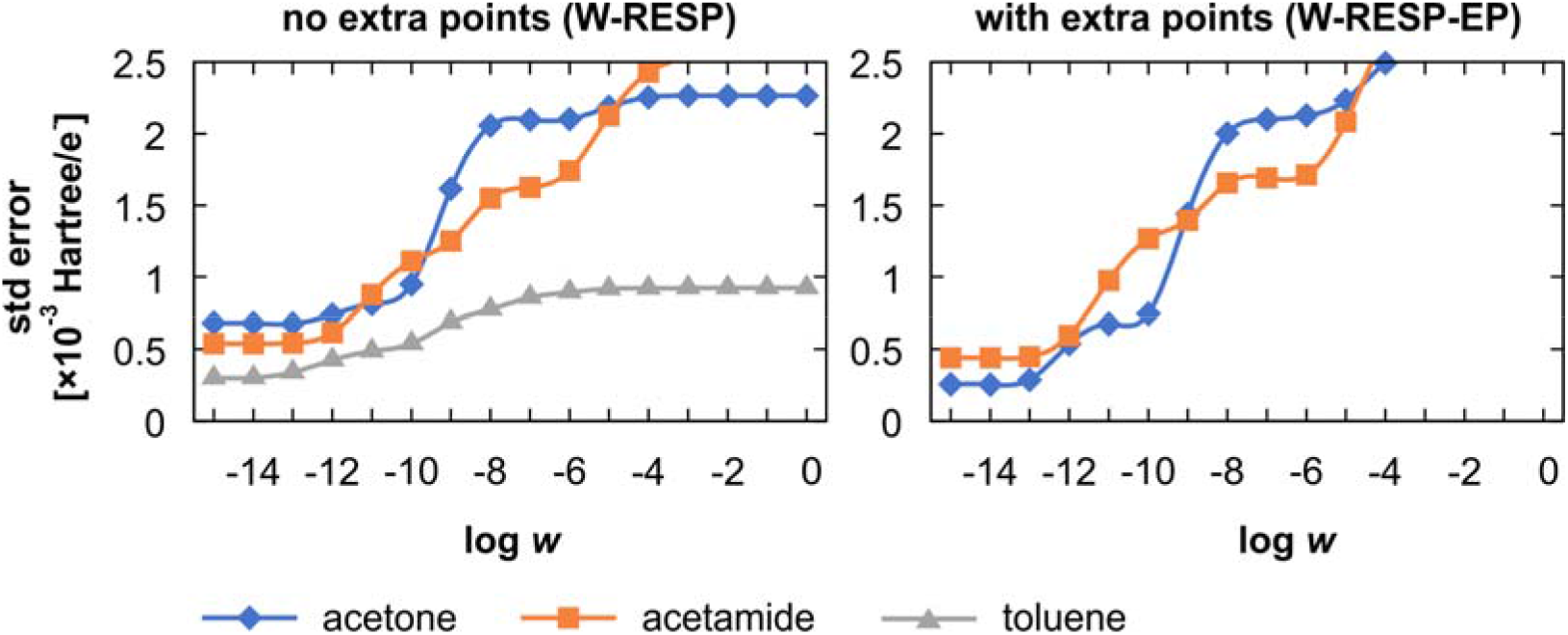
Effect of restraint scaling coefficient *w* on the overall quality of fit represented by the SE (defined in equation 5) for acetone, acetamide and toluene molecules. The left panel corresponds to W-RESP fits without extra point charges, whereas the right panel corresponds to W-RESP-EP fits (note that toluene did not have any extra point charge).

**Table 4:** Comparison of the partial charges of acetone, accuracy of corresponding ESP potential on a W-RESP grid represented by the SE (in ×10^−3^ Hartree/e^−^, see equation 5), and dipole and quadrupole moments (in debye and debye·angstroms, respectively) for ESP, W-RESP, W-RESP-EP and CM5 charges. For comparison, the ESP charges were calculated with equivalencing in a two stage model.

| atom | without EPs |  |  | with EPs |  |  | QM |
| --- | --- | --- | --- | --- | --- | --- | --- |
|  | ESP | W-RESP | CM5 | ESP | W-RESP-EP | CM5 |  |
| Charges |  |  |  |  |  |  |  |
| C(methyl) | -0.602 | -0.520 | -0.247 | -0.643 | -0.537 | -0.247 |  |
| H | 0.168 | 0.148 | 0.110 | 0.175 | 0.149 | 0.110 |  |
| C(carbonyl) | 0.724 | 0.675 | 0.172 | 0.816 | 0.747 | 0.172 |  |
| O | -0.529 | -0.523 | -0.332 | 0.281 | 0.213 | 0.363 |  |
| EP | - | - | - | -0.430 | -0.390 | -0.347 |  |
| Standard Error |  |  |  |  |  |  |  |
|  | 0.682 | 0.677 | 2.266 | 0.257 | 0.287 | 2.590 |  |
| Dipole and quadrupole moments |  |  |  |  |  |  |  |
| $\mu$ | 3.417 | 3.420 | 3.288 | 3.470 | 3.469 | 3.186 | 3.468 |
| $Q_{xx}$ | 5.948 | 5.977 | 7.324 | 4.343 | 4.496 | 5.971 | 1.878 |
| $Q_{yy}$ | 2.717 | 2.675 | -1.153 | 3.282 | 3.154 | -1.290 | 1.367 |
| $Q_{zz}$ | -8.665 | -8.652 | -6.171 | -7.624 | -7.651 | -4.680 | -3.245 |

### Test case 1: AT, AU and GC base pairing

The performance of the newly developed W-RESP and W-RESP-EP charges was assessed by calculating the gas-phase interaction pairing energies of AT, AU and GC base pairs. The standard HF/6-31G(d) RESP charges^39–40^ are known to significantly underestimate interaction energies compared to QM benchmarks^73^, which may contribute to inaccuracies in simulations of nucleic acids with the current state-of-the-art RNA *ff*s^91^. The underestimated strength of base pairing in MD simulations results in, e.g., perturbation of helical stems due to excessive and irreversible base pair fraying at the helix termini.^92^ It can also compromise enhanced sampling folding simulations, as the stability of canonical A-RNA stems may be underestimated.^43, 50, 93–94^ Recently, we introduced an additional force field term called gHBfix^50^ that can be used to tune non-bonded interactions in the force field, such as hydrogen bonding. We showed that the gHBfix term might improve the performance of the RNA force field by stabilization of underestimated base pairing interactions. Here, we show that inaccurate description of the charge model might explain at least part of the observed base pairing understabilization. Obviously, the stability of base pairing in MD simulations will also be affected by the chosen water model and lack of polarization.

We benchmarked the MM base pairing energies calculated with RESP, ^39–40^ RESP2,^61, 95^ W-RESP and W-RESP-EP charges against high-level QM data. All calculations were performed in the gas phase using rigid planar nucleobases. The reference high-level QM energies were calculated at CBS(T) level (see Methods). Six different base pair stretch values were used for each AU, AT and GC base pair (see Methods). The W-RESP charge revealed a significant improvement of the base pairing energies only for the GC base pair, whereas for the AU and AT base pairs, the accuracy of the base pairing energies was comparable to that of the RESP charge set Figures 5). The best results were achieved using the W-RESP-EP charge set with EPs Figures 5), which performed well for all three base pairs. Notably, the W-RESP-EP charges were able to correctly predict optimal stretch value for all three base pairs, whereas the standard AMBER force field using RESP charges tended to overestimate the base pair stretch by ~0.1 Å. We propose that additional tuning of the MM base pairing interaction energies at the top of the W-RESP-EP charge model might be achieved by further adjustment of the van der Waals terms. However, this was beyond the scope of the present study.

**Figure 5:**
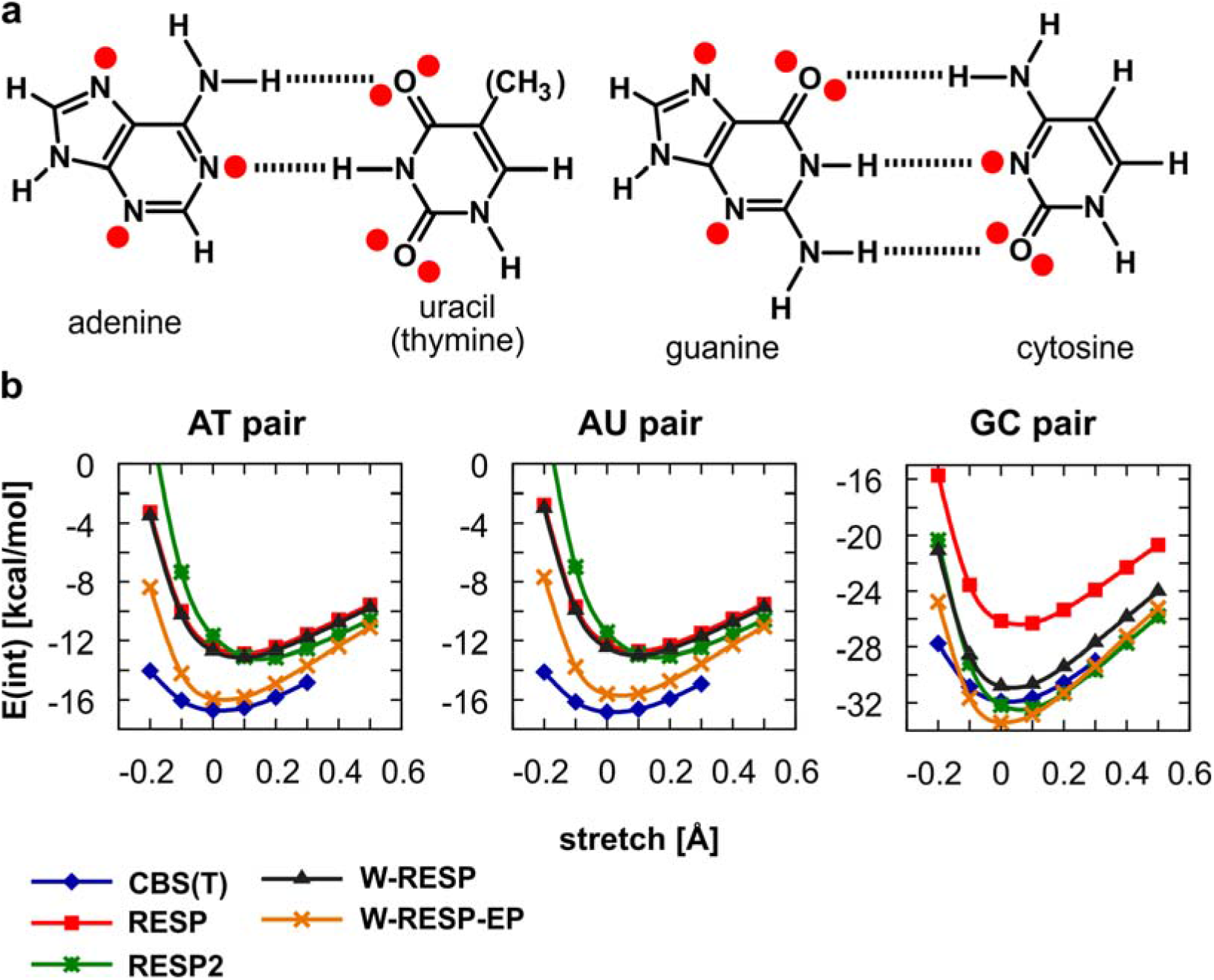
(a) Schemes for AU, AT and GC base pairs with hydrogen bonds (dashed lines) with highlighted extra point charges (red dots) (b) QM CBS(T) and MM base-pairing energies between rigid planar nucleobases calculated for AT, AU and GC base pairs. The MM energies were calculated with standard RESP charges, recently published RESP2 charges, and the charge models W-RESP and W-RESP-EP presented in this study. All MM energies were calculated using Amber Lennard-Jones parameters except for RESP2 charges, which were used along with recommended Lennard-Jones parameters^61^.

### Test case 2: MD simulations of A-RNA duplex

The W-RESP and W-RESP-EP charges were next tested on two canonical A-RNA duplexes, i.e., the r(GCACCGUUGG)_2_ decamer (10-mer) and r(UUAUAUAUAUAUAA)_2_ tetradecamer (14-mer). We performed four MD simulations in total, i.e., two standard MD simulations for each duplex, one with the W-RESP and the other with the W-RESP-EP charge potential. We inspected the overall stability, fluctuation of helical parameters and base-pair fraying at the end of both duplexes. The results were compared with simulations using the same RNA *ff* (see Methods) with standard RESP charges and with our recently suggested correction for RNA simulations using the external gHBfix potential.^50^

**Table MD1:** Analysis of base-pair fraying of 10-mer and 14-mer A-RNA test duplexes from 1 μs-long MD simulations with standard RESP charges (control), standard charges and the external gHBfix potential (gHBfix),^50^ W-RESP charges and W-RESP-EP charges. The frequency of fraying was estimated by calculating the root mean square deviation (RMSD, considering nucleobase atoms of each terminal base pair) with a RMSD cutoff 1.35 Å.

|  | top [%]<sup>a</sup> |  |  |  | bottom [%]<sup>b</sup> |  |  |  |
| --- | --- | --- | --- | --- | --- | --- | --- | --- |
|  | control<sup>c</sup> | gHBfix<sup>d</sup> | W-RESP<sup>e</sup> | W-RESP-EP<sup>f</sup> | control<sup>c</sup> | gHBfix<sup>d</sup> | W-RESP<sup>e</sup> | W-RESP-EP<sup>f</sup> |
| 10-mer | 0.2 | 0.0 | 0.1 | 0.1 | 0.5 | 0.0 | 0.2 | 0.0 |
| 14-mer | 14.6 | 1.5 | 7.1 | 1.7 | 47.3 | 1.3 | 16.3 | 7.7 |

<sup>a</sup> Population of frayed states for the top base pair, i.e., nucleotides 1-20 and 1-28 for the decamer and tetradecamer, respectively.
<sup>b</sup> Population of frayed states for the bottom base pair, i.e., nucleotides 10-11 and 14-15 for the decamer and tetradecamer, respectively.
<sup>c</sup> RNA ff (see Methods) with standard RESP<sup>78</sup> charges. Data taken from Ref. <sup>50</sup>.
<sup>d</sup> Same ff as in <sup>c</sup> with external gHBfix potential. Data taken from Ref. <sup>50</sup>.
<sup>e</sup> Same ff as in <sup>c</sup> with W-RESP charge parameterization introduced here (see Methods).
<sup>f</sup> Same ff as in <sup>c</sup> with W-RESP-EP charge parameterization introduced here (see Methods).

All four simulations with our novel charge models revealed stable behavior with the RMSD of all heavy atoms from the starting X-ray structure fluctuating around 1.24 ± 1.28 Å (W-RESP; 1.17 ± 1.21 Å with the W-RESP-EP charge model) and 1.91 ± 1.96 Å (W-RESP; 1.88 ± 1.93 Å with the W-RESP-EP charge model) for the 10-mer and 14-mer, respectively. These RMSD values are comparable to our previous results from A-RNA simulations with the standard *ff*99bsc0χ_OL3_^35, 77–79^ RNA *ff* (see, e.g., Refs. ^50, 96^ for details). Base-pair breathing (fraying) of GC pairs at the end of helices occurred rarely (in both the W-RESP and W-RESP-EP simulations), whereas AU base-pair fraying was more frequent (Table MD1). However, all identified opened states of AU base pairs were able to reform back into the canonical Watson-Crick base pair and the overall frequency of fraying was still significantly reduced in comparison with standard *ff*99bsc0χ_OL3_ simulations.^50, 92^ Note that accurate (unambiguous) experimental data for the quantification of frayed structures are not available for RNA sequences.^92^ Recent MD simulations have shown that base-pair openings are usually followed by formation of likely spurious long-lived noncanonical interactions.^92^ In other words, end fraying has a tendency to propagate further along the RNA helix, which most likely is a simulation artifact. It is known, e.g., to detrimentally affect simulations of some protein-RNA complexes.^97^ Thus, the identified reduction of base-pair fraying in the simulations with both the W-RESP and W-RESP-EP charge models is expected to have positive effects, mimicking the improvement achieved by the gHBfix potential.^50^ This may be important because an improved charge model would be a preferred solution over the gHBfix correction. In other words, the better the performance achieved by basic RNA force-field terms, the more flexibility there is for further tuning by the gHBfix potentials.

**Table MD2:**
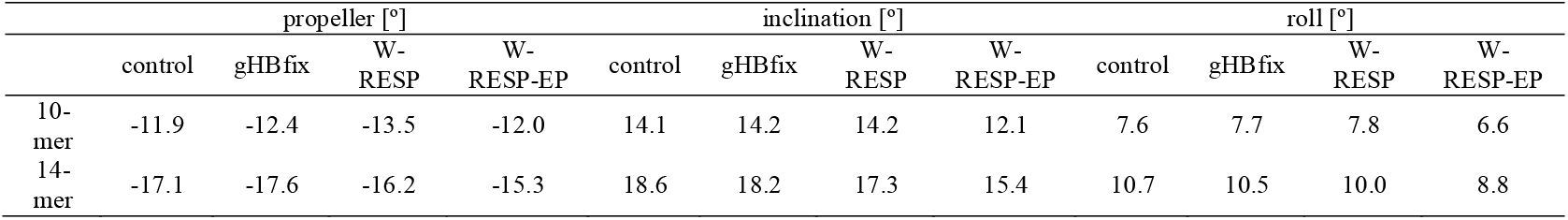
Analysis of helical base-pair parameters for two simulated duplexes, i.e., the r(GCACCGUUGG)_2_ decamer (10-mer) and r(UUAUAUAUAUAUAA)_2_ tetradecamer (14-mer). Averaged values of base-pair propeller, roll and inclination from 1 μs-long MD simulations with various settings (see Table MD1 for details). Helical parameters were obtained by CPPTRAJ^98^ and calculated for the inner six and ten base-pair segments of the 10-mer and 14-mer, respectively.

|  | propeller [°] |  |  |  | inclination [°] |  |  |  | roll [°] |  |  |  |
| --- | --- | --- | --- | --- | --- | --- | --- | --- | --- | --- | --- | --- |
|  | control | gHBfix | W-RESP | W-RESP-EP | control | gHBfix | W-RESP | W-RESP-EP | control | gHBfix | W-RESP | W-RESP-EP |
| 10-mer | -11.9 | -12.4 | -13.5 | -12.0 | 14.1 | 14.2 | 14.2 | 12.1 | 7.6 | 7.7 | 7.8 | 6.6 |
| 14-mer | -17.1 | -17.6 | -16.2 | -15.3 | 18.6 | 18.2 | 17.3 | 15.4 | 10.7 | 10.5 | 10.0 | 8.8 |

We subsequently analyzed the helical base-pair parameters, i.e., those measuring orientation and displacement of Watson–Crick base pairs. Application of the W-RESP and W-RESP-EP charge potentials did not significantly affect the helical parameters of the simulated A-RNA duplexes. The average values of base-pair propeller, inclination and roll (main descriptors of A-RNA duplexes^96, 99^) fluctuated near values observed within a control simulation with the standard RESP charge model (Table MD2). Thus, the novel W-RESP and W-RESP-EP charge models revealed stable simulations of A-RNA duplexes where helical parameters were minimally affected, whereas the terminal base pairs of A-RNA duplexes were evidently stabilized, at least kinetically, but most likely also thermodynamically.

## CONCLUSIONS

The electrostatic term is a crucial component of the standard non-polarizable force fields. The conformational dynamics of biomolecules is significantly influenced by nonbonded interactions, including solvation by explicit waters. As we have argued elsewhere, the van der Waals parameters affect the interaction energy indirectly via electrostatics, and thus their potential for fine-tuning of the force field performance is rather limited.^50^ On the other hand, even small changes in the parametrization of the electrostatic term may significantly influence the overall conformational preference of the simulated system, in particular for highly charged systems, such as nucleic acids. In this study, we analyzed the standard RESP charge derivation method. The broad success of the RESP method is likely because the RESP procedure parameterizes the partial charges to reproduce the QM ESP around the molecule. Thus, the molecules interact with each other via an electrostatic field that as closely as possible approaches the QM quality. However, the quality of the RESP fit, i.e., residual discrepancy between the ESP potential generated by RESP partial charges and that calculated at the QM level, varied among different types of molecules. The worst ability of RESP charges to reproduce the QM ESP potential was observed in aromatic heterocyclic molecules, including nucleobases of RNA and DNA. This suggests that the parameterization of the partial charges in AMBER nucleic acid force fields might be of lower quality compared to, e.g., AMBER protein force fields. Indeed, we recently reported that limitations in the description of nonbonded terms, in particular underestimated base pairing and overestimated sugar-phosphate interactions, are responsible for difficulties in describing the folding free energy landscape of small RNA oligonucleotides.^43, 50, 100^ Underestimation of base paring interactions has also been reported in DNA^73^ and suggested to be responsible for extensively overestimated base pair fraying in DNA.^101^ Here, we showed that the proposed charge derivation model, in particular the version with EP charges W-RESP-EP, is able to eliminate the underestimation of the base pairing and significantly improve the behavior of terminal base pairs in MD simulations. Nonetheless, careful and extensive testing of the force field with reparametrized W-RESP(-EP) charges is required to fully understand their performance. Such extensive testing, which was beyond the scope of this study, is under progress in our lab and will be addressed separately.

Finally, it is worth noting that we applied the W-RESP(-EP) charges in the RNA force field only to atoms of nucleobases, whereas partial charges of the atoms along the sugar-phosphate backbone were kept intact. The reparametrization of the nucleobase charges required only minimal additional adjustment of other force field terms as nucleobases are rather rigid and the only torsion term affected by the new partial charges corresponded to glycosidic bonds. Complete reparametrization of the partial charges in nucleic acid force fields, including the charges of sugar-phosphate backbone atoms, would require extensive reparametrization of all dihedral parameters and likely also the van der Waals terms. Therefore, we suggest the need for careful testing of the potential performance improvement in the hybrid force field with W-RESP(-EP) charges on nucleobase atoms and RESP on the sugar-phosphate backbone prior to extensive reparametrization of the complete force field.

## Supporting information

Supporting Information for the article

## Acknowledgements

PB, PK, and MJ were supported by Czech Science Foundation grant No 18-25349S. JS and VM were supported by Czech Science Foundation grant No 20-16554S.

