## Supporting Information for the article for "W-RESP: Well-Restrained Electrostatic Potential Derived Charges. Revisiting the Charge Derivation Model"

### Atomic radii for the grid definition

**Table S1** - Standard error of the ESP fit (as defined in equation 5 in the Main text), minimal (min) and maximal (max) difference between MM and QM ESP potential at the grid point $\widehat{V_i} - V_i$ (see equation 1 in the Main text) in $\times 10^{-3}$ Hartree/$e^-$, all calculated on the same W-RESP grid. Compounds are sorted from largest to lowest reduction of the standard error.

| | MK radii | | | W-RESP radii | | | |
|---|---|---|---|---|---|---|---|
| | **Std error** | **Min** $\widehat{V_i} - V_i$ | **Max** $\widehat{V_i} - V_i$ | **Std error** | **Min** $\widehat{V_i} - V_i$ | **Max** $\widehat{V_i} - V_i$ | **Reduction of std error [%]** |
| toluene | 0.387 | -2.091 | 2.082 | 0.301 | -2.930 | 1.320 | 22.3 |
| indole | 0.411 | -4.029 | 1.880 | 0.328 | -3.869 | 1.850 | 20.2 |
| methane | 0.170 | -1.022 | 0.583 | 0.138 | -0.947 | 0.431 | 19.2 |
| benzene | 0.497 | -2.179 | 1.329 | 0.403 | -3.543 | 1.116 | 18.9 |
| 3-methylindole | 0.433 | -2.819 | 2.403 | 0.355 | -2.720 | 2.180 | 18.1 |
| methyluracil | 0.618 | -7.244 | 3.978 | 0.544 | -5.202 | 4.734 | 12.0 |
| thymine | 0.677 | -7.595 | 4.587 | 0.597 | -5.481 | 5.076 | 11.8 |
| uracil | 0.753 | -7.559 | 4.368 | 0.665 | -5.516 | 4.964 | 11.7 |
| acetamide | 0.609 | -7.921 | 3.865 | 0.541 | -6.242 | 4.271 | 11.2 |
| methylcytosine | 0.571 | -7.673 | 3.324 | 0.507 | -5.983 | 2.946 | 11.2 |
| cytosine | 0.752 | -9.337 | 5.288 | 0.682 | -7.206 | 6.138 | 9.3 |
| guanidinium | 0.330 | -2.819 | 0.996 | 0.302 | -2.427 | 1.057 | 8.7 |
| methylphosphate | 1.189 | -7.956 | 5.336 | 1.090 | -8.368 | 4.679 | 8.4 |
| methylguanine | 0.816 | -7.701 | 8.096 | 0.750 | -8.663 | 5.722 | 8.1 |
| guanine | 0.837 | -7.691 | 7.719 | 0.777 | -8.438 | 6.199 | 7.3 |
| acetate | 1.179 | -10.068 | 5.365 | 1.101 | -7.903 | 5.416 | 6.7 |
| propionate | 1.192 | -10.136 | 8.733 | 1.123 | -7.711 | 9.073 | 5.8 |
| dimethylphosphate | 1.035 | -7.380 | 5.339 | 0.980 | -7.447 | 4.151 | 5.3 |
| propionic acid | 0.754 | -6.609 | 5.579 | 0.715 | -5.538 | 5.791 | 5.3 |
| N-methylacetamide | 0.967 | -8.134 | 5.713 | 0.920 | -6.323 | 6.612 | 4.9 |
| imidazole | 1.214 | -5.846 | 12.264 | 1.156 | -6.687 | 10.046 | 4.8 |
| methyladenine | 0.823 | -8.220 | 8.227 | 0.792 | -8.515 | 6.496 | 3.7 |
| adenine | 0.845 | -7.839 | 7.644 | 0.817 | -8.460 | 5.985 | 3.3 |
| methylammonium | 0.879 | -3.625 | 5.597 | 0.852 | -3.243 | 4.685 | 3.1 |
| ethanethiol | 2.047 | -14.750 | 12.960 | 1.989 | -15.194 | 12.334 | 2.8 |
| ethylammonium | 0.697 | -2.534 | 5.088 | 0.678 | -2.826 | 5.391 | 2.7 |
| acetic acid | 0.827 | -6.476 | 5.901 | 0.808 | -5.329 | 6.572 | 2.2 |
| propionylamide | 0.713 | -5.440 | 4.062 | 0.702 | -4.609 | 4.670 | 1.4 |
| ethane | 0.770 | -2.790 | 3.192 | 0.759 | -2.965 | 3.166 | 1.4 |
| propane | 0.649 | -3.625 | 3.866 | 0.640 | -3.761 | 4.013 | 1.3 |
| ADNA_sugar | 1.027 | -7.346 | 7.088 | 1.015 | -7.530 | 6.957 | 1.1 |
| dimethylether | 1.068 | -4.199 | 8.064 | 1.057 | -4.119 | 7.616 | 1.0 |
| dimethylsulfide | 1.766 | -13.955 | 12.569 | 1.749 | -14.586 | 12.056 | 1.0 |
| B-DNA sugar | 1.149 | -6.840 | 5.994 | 1.141 | -7.780 | 5.597 | 0.7 |
| phenol | 1.161 | -7.453 | 4.257 | 1.153 | -7.247 | 4.198 | 0.7 |
| ethanethiol cf2 | 2.789 | -22.281 | 14.794 | 2.772 | -23.035 | 14.496 | 0.6 |

| | | | | | | | |
|---|---|---|---|---|---|---|---|
| ethylamine | 1.179 | -7.250 | 6.375 | 1.174 | -7.187 | 6.641 | 0.4 |
| ethanol | 0.843 | -4.824 | 5.864 | 0.845 | -4.609 | 5.567 | -0.3 |
| dimethylamine | 1.965 | -13.729 | 4.747 | 1.971 | -14.429 | 5.015 | -0.3 |
| methanethiol | 3.015 | -24.363 | 13.106 | 3.025 | -25.259 | 12.801 | -0.3 |
| isopropanol | 1.140 | -7.556 | 5.832 | 1.154 | -7.813 | 6.084 | -1.3 |
| isopropanol cf2 | 0.557 | -6.641 | 3.866 | 0.565 | -5.433 | 3.119 | -1.6 |
| methylguanidine | 1.166 | -7.605 | 4.870 | 1.189 | -7.757 | 5.385 | -2.0 |
| ethanol cf2 | 1.572 | -8.940 | 7.531 | 1.608 | -9.228 | 7.403 | -2.3 |
| ethylamine cf2 | 1.534 | -9.479 | 6.022 | 1.576 | -9.726 | 6.161 | -2.8 |
| 4-methylimidazole | 0.940 | -5.117 | 10.191 | 0.972 | -6.685 | 7.326 | -3.4 |
| acetone | 0.659 | -7.988 | 5.637 | 0.682 | -7.118 | 6.213 | -3.5 |
| methylamine | 1.948 | -9.659 | 6.102 | 2.026 | -9.770 | 6.276 | -4.0 |
| methylthymine | 0.534 | -6.382 | 4.344 | 0.558 | -4.823 | 4.887 | -4.4 |
| methanol | 1.660 | -10.124 | 6.041 | 1.756 | -10.664 | 5.623 | -5.8 |
| A-RNA sugar | 1.716 | -10.616 | 10.240 | 1.834 | -11.458 | 10.826 | -6.9 |
| isobutane | 0.346 | -2.561 | 1.202 | 0.433 | -3.033 | 1.148 | -25.0 |

### Extra-points

To obtain the robust parameters possible final position of an extra point charge (EP) was determined as consensus amongst all compounds from our training set carrying the particular functional group. For most functional groups carrying N or O atoms we found that absolute values of charge magnitude of EP or associated heavy atom significantly increase above 1 a.u. under the atom-EP distance of 0.25Å (Figures S1C, S2C, S3D and S7D) which makes it the lower border for a consensus position of the EPs. Furthermore, improvement of the standard error by the EPs usually does not significantly depend on the atom-EP distance (see Figures S1B, S2B, S3C, S7C, S8B, S9C and S11C). Thus, theoretically any distance from explored range of 0.25 up to 0.5Å can be used as the consensus atom-EP distance. Therefore, we used the distance of 0.35Å suggested by study of Dixon et al.[1]

Analogic trends are valid also for compounds carrying sulfur atom: The absolute values of charge magnitude of EP or sulfur significantly increase above 1 a.u. under the S-EP distance of 0.6Å (Figures S5D, S10D,E) and again, the improvement of the standard error of the ESP fit does not significantly change with the S-EP distance (Figures S5C, S10C). Thus, we used the S-EP distance of 0.7Å as did refs[1-2].

*Imine group*

One EP located in intuitive position of the electron lone pair already proposed by refs.[1, 3-4] (Figure S1A) significantly improves the standard error of the ESP fit for all appropriate compounds from our training set (Figure S1B, Table S2). Note that the least improved standard error was in case of guanine and cytosine (and their methyl derivatives) because they carry also carbonyl group which requires another EPs (Table S2 – compounds marked “(CO)”).

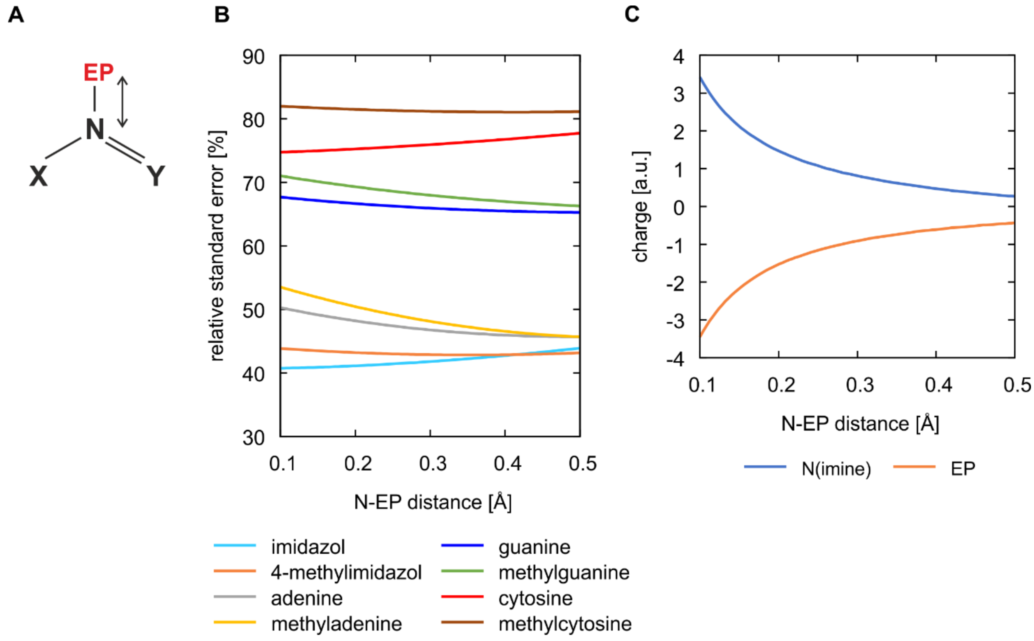

**Figure S1** – **A)** The EP charge is located on bisector of X-N-Y outer angle whereas various N-EP distances are probed. **B)** Relative standard error (with respect to model without EPs) of the ESP fit equivalenced in 2 stages including 1 EP on each imine group as it depends on N-EP distance, of all compounds with imine groups from our training set. **C)** Charge magnitudes of imine nitrogen and EP of imidazole as they depend on N-EP distance.

**Table S2 -** Impact of including 1 EP on nitrogen atom of each imine group (with sp2-like position and N-EP distance of 0.35Å – see Figure 3 in the Main text) on standard error of the ESP fit equivalenced in 2 stages and on minimal (min) and maximal (max) $V_i^{MM}$-$V_i^{QM}$ (all: ×$10^{-3}$ Hartree/$e^-$).

| | **no EPs on imine** | | | **1 EP per imine** | | | **Relative Std Error (EP: no EP) [%]** |
|---|---|---|---|---|---|---|---|
| | **Std Error** | **Min $V_i^{MM}$-$V_i^{QM}$** | **Max $V_i^{MM}$-$V_i^{QM}$** | **Std Error** | **Min $V_i^{MM}$-$V_i^{QM}$** | **Max $V_i^{MM}$-$V_i^{QM}$** | |
| **imidazole** | 1.156 | -6.687 | 10.046 | 0.488 | -4.259 | 1.887 | 42.2 |
| **4-methylimidazole** | 0.972 | -6.685 | 7.326 | 0.416 | -4.106 | 1.988 | 42.8 |
| **adenine** | 0.817 | -8.460 | 5.985 | 0.378 | -4.280 | 1.739 | 46.3 |
| **methyladenine** | 0.792 | -8.515 | 6.496 | 0.374 | -4.637 | 1.537 | 47.2 |
| **guanine** | 0.777 | -8.438 | 6.199 | 0.510 | -5.055 | 5.561 | 65.7 |
| **guanine (CO)** | 0.719 | -7.717 | 6.249 | 0.428 | -4.827 | 1.894 | 59.5 |
| **methylguanine** | 0.750 | -8.663 | 5.722 | 0.506 | -5.309 | 5.518 | 67.4 |
| **methylguanine (CO)** | 0.694 | -7.951 | 6.536 | 0.421 | -4.728 | 1.798 | 60.6 |
| **cytosine** | 0.682 | -7.206 | 6.138 | 0.521 | -5.463 | 5.784 | 76.3 |
| **cytosine (CO)** | 0.498 | -4.758 | 3.493 | 0.376 | -3.308 | 1.494 | 75.4 |
| **methylcytosine** | 0.507 | -5.983 | 2.946 | 0.411 | -4.824 | 2.865 | 81.1 |
| **methylcytosine (CO)** | 0.406 | -4.343 | 3.405 | 0.341 | -3.164 | 1.417 | 84.0 |
| **imidazole** | 1.156 | -6.687 | 10.046 | 0.488 | -4.259 | 1.887 | 42.2 |

**(CO)=including 2 EPs on carbonyl group**

*Carbonyl group*

The optimal position of two symmetric EPs has C-O-EP angle ranging between 60 and 90 degrees (depends on compound and also on the O-EP distance; Figures S2 A,B) which is in agreement with ref. [3, 5]. At the distance of 0.35Å the consensus optimal C-O-EP angle is 85 degrees (Figure S2A, Table S3).

Two EPs in intuitive positions of the electron lone pairs suggested by Dixon et al. [1] only marginally improve the ESP fit (Figure S2A, Table S4) and are thus unsuitable for our purpose.

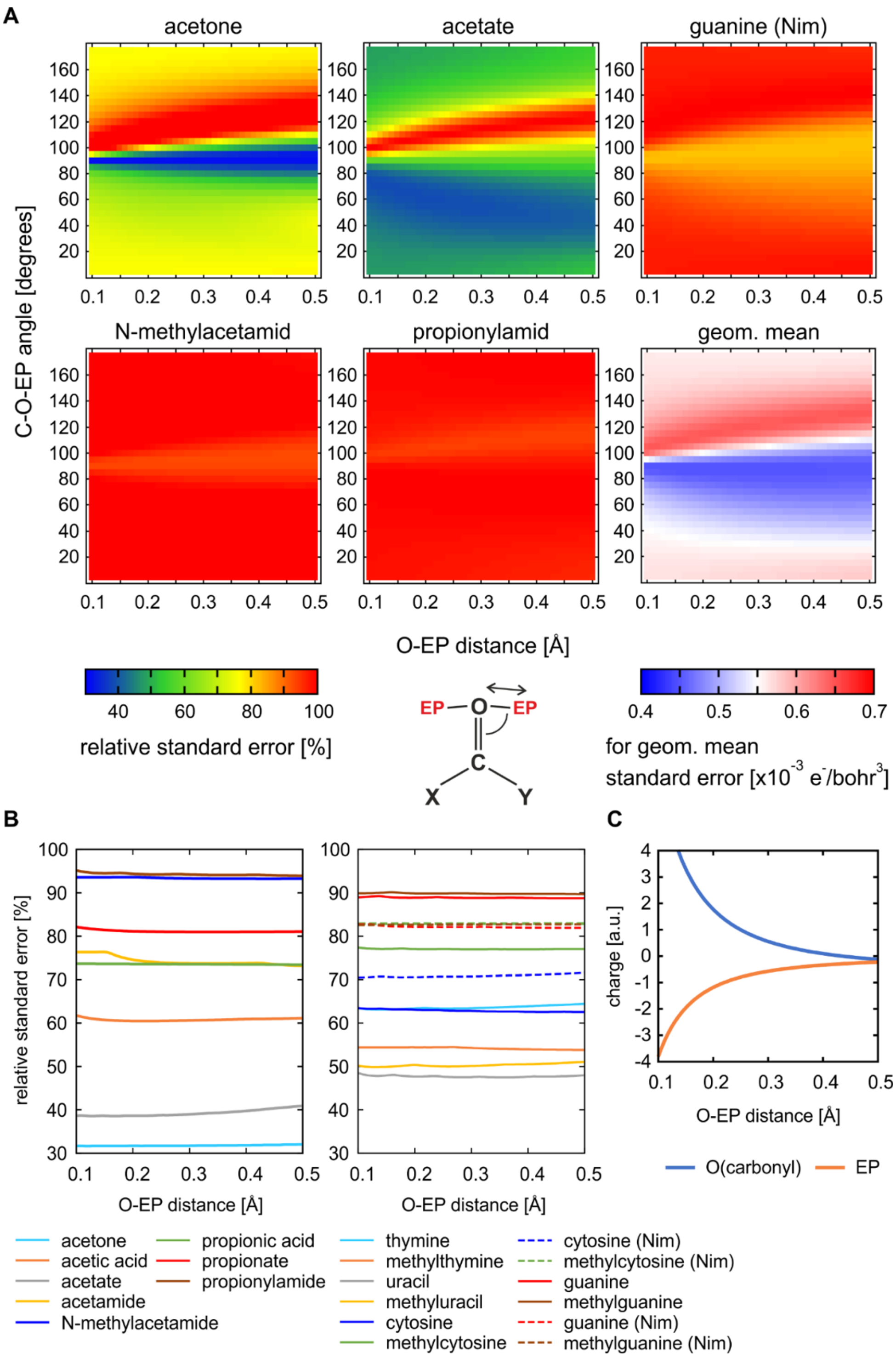

**Figure S2** – **A**) Impact of EPs position (O-EP distance and C-O-EP angle in the plane of the CO group) on the relative standard error (relative to the one without EPs) of ESP fit equivalenced in 2 stages of selected compounds shown on 2D maps and geometric mean of the standard error [x10$^{-3}$ Hartree/e$^{-}$] of all compounds with carbonyl group from the training set **B)** How relative standard error depends on O-EP distance (C-O-EP angle is always optimal for the given distance) **C**) Charge magnitudes of carbonyl oxygen and EP of acetone as they depend on O-EP distance at C-O-EP angle of 85°.

**Table S3** – Impact of including 2 symmetric EPs on oxygen atom of carbonyl group (O-EP distance=0.35Å, C-O-EP angle=85°, EPs lie at the plane of the carbonyl group - see Figure 3 in the Main text) on standard error of the ESP fit equivalenced in 2 stages and on minimal and maximal $V_i^{MM}$-$V_i^{QM}$ (all: ×$10^{-3}$ Hartree/$e^-$).

| | **no EPs on carbonyl** | | | **2 EPs per carbonyl (ours, angle=85°)** | | | **Relative Std Error** |
|---|---|---|---|---|---|---|---|
| | **Std Error** | **Min $V_i^{MM}$-$V_i^{QM}$** | **Max $V_i^{MM}$-$V_i^{QM}$** | **Std Error** | **Min $V_i^{MM}$-$V_i^{QM}$** | **Max $V_i^{MM}$-$V_i^{QM}$** | **(EP:no EP) [%]** |
| **acetone** | 0.682 | -7.118 | 6.213 | 0.257 | -3.150 | 0.909 | 37.7 |
| **acetic acid** | 0.808 | -5.329 | 6.572 | 0.491 | -4.534 | 2.061 | 60.7 |
| **acetic acid (OH)** | 0.820 | -5.298 | 6.544 | 0.494 | -5.171 | 1.886 | 60.3 |
| **acetate** | 1.101 | -7.903 | 5.416 | 0.585 | -3.857 | 1.744 | 53.2 |
| **acetamide** | 0.541 | -6.242 | 4.271 | 0.440 | -3.221 | 1.464 | 81.5 |
| **N-methylacetamide** | 0.920 | -6.323 | 6.612 | 0.874 | -5.317 | 3.722 | 95.0 |
| **propionic acid** | 0.715 | -5.538 | 5.791 | 0.529 | -5.285 | 3.330 | 74.0 |
| **propionic acid (OH)** | 0.664 | -5.444 | 5.691 | 0.504 | -3.981 | 3.593 | 75.9 |
| **propionate** | 1.123 | -7.711 | 9.073 | 0.909 | -5.012 | 4.385 | 81.0 |
| **propionylamide** | 0.702 | -4.609 | 4.670 | 0.699 | -3.826 | 4.169 | 99.5 |
| **thymine** | 0.597 | -5.481 | 5.076 | 0.384 | -3.830 | 2.078 | 64.3 |
| **methylthymine** | 0.558 | -4.823 | 4.887 | 0.313 | -2.545 | 2.777 | 56.0 |
| **uracil** | 0.665 | -5.516 | 4.964 | 0.376 | -2.976 | 1.125 | 56.5 |
| **methyluracil** | 0.544 | -5.202 | 4.734 | 0.283 | -2.526 | 1.105 | 51.9 |
| **cytosine** | 0.682 | -7.206 | 6.138 | 0.498 | -4.758 | 3.493 | 73.1 |
| **cytosine (Nim)** | 0.521 | -5.463 | 5.784 | 0.376 | -3.308 | 1.494 | 72.2 |
| **methylcytosine** | 0.507 | -5.983 | 2.946 | 0.406 | -4.343 | 3.405 | 80.1 |
| **methylcytosine (Nim)** | 0.411 | -4.824 | 2.865 | 0.341 | -3.164 | 1.417 | 83.0 |
| **guanine** | 0.777 | -8.438 | 6.199 | 0.719 | -7.717 | 6.249 | 92.6 |
| **guanine (Nim)** | 0.510 | -5.055 | 5.561 | 0.428 | -4.827 | 1.894 | 83.9 |
| **methylguanine** | 0.750 | -8.663 | 5.722 | 0.694 | -7.951 | 6.536 | 92.5 |
| **methylguanine (Nim)** | 0.506 | -5.309 | 5.518 | 0.421 | -4.728 | 1.798 | 83.1 |

**(OH)=including 1 EP on hydroxyl group (see Figure 3 in the Main text)**
**(Nim)=including 1 EP on each imine group (see Figure 3 in the Main text)**

**Table S4** – Impact of including 2 symmetric EPs on oxygen atom of carbonyl group placed at position suggested by Dixon et al.[1] (O-EP distance=0.35Å, C-O-EP angle=120°, EPs lie at the plane of the carbonyl group) on standard error of the ESP fit equivalenced in 2 stages and on minimal and maximal $V_i^{MM}$-$V_i^{QM}$ (all: ×$10^{-3}$ Hartree/$e^-$).

| | 2 EPs per carbonyl (Dixon, angle=120°) | | | Relative Std Error (EP:no EP) [%] |
|---|---|---|---|---|
| | Std Error | Min $V_i^{MM}$-$V_i^{QM}$ | Max $V_i^{MM}$-$V_i^{QM}$ | |
| **acetone** | 0.751 | -6.392 | 5.418 | 110.0 |
| **acetic acid** | 0.823 | -5.534 | 6.165 | 101.8 |
| **acetic acid (OH)** | 0.846 | -5.796 | 5.988 | 102.5 |
| **acetate** | 1.079 | -7.439 | 5.778 | 98.1 |
| **acetamide** | 0.541 | -6.167 | 4.316 | 100.1 |
| **N-methylacetamide** | 0.933 | -6.157 | 7.257 | 101.4 |
| **propionic acid** | 0.697 | -6.045 | 5.091 | 97.6 |
| **propionic acid (OH)** | 0.698 | -6.024 | 5.099 | 97.6 |
| **propionate** | 1.142 | -8.205 | 8.197 | 101.7 |
| **propionylamide** | 0.671 | -5.428 | 4.181 | 95.6 |
| **thymine** | 0.584 | -5.687 | 4.403 | 97.7 |
| **methylthymine** | 0.500 | -5.172 | 3.505 | 89.7 |
| **uracil** | 0.662 | -5.540 | 4.778 | 99.6 |
| **methyluracil** | 0.533 | -5.658 | 3.864 | 98.0 |
| **cytosine** | 0.673 | -6.630 | 6.886 | 98.8 |
| **cytosine (Nim)** | 0.511 | -5.858 | 4.945 | 98.2 |
| **methylcytosine** | 0.505 | -5.767 | 3.193 | 99.7 |
| **methylcytosine (Nim)** | 0.397 | -5.419 | 2.147 | 96.7 |
| **guanine** | 0.717 | -7.785 | 6.447 | 92.3 |
| **guanine (Nim)** | 0.475 | -5.720 | 3.608 | 93.2 |
| **methylguanine** | 0.703 | -8.032 | 5.830 | 93.7 |
| **methylguanine (Nim)** | 0.484 | -5.887 | 4.012 | 95.6 |

**(OH)=including 2 EPs on hydroxyl group as suggested by Dixon et al.[1]**
**(Nim)=including 1 EP on each imine group as suggested by Dixon et al.[1]**

*Hydroxyl*

According to our data, 1 EP (located in the C-O-H plane with C-O-EP angle of 290°, see Figure 3 in the Main text) provides satisfactory improvement of the ESP fit of aliphatic alcohols, as well as of ribose in A-RNA-like conformation and deoxyribose in both A- and B-DNA-like conformations (Figures S3A,B, Table S5). However, its contribution to the improvement of acetic and propionic acids is minor, in these cases the major improvement is provided by EPs on carbonyl group.

Similarly to the carbonyl functional group, the position of EPs suggested by Dixon et al.[1] (with intuitive sp3-like geometry on oxygen) was not designed with the aim to improve the ESP fit and is unsuitable for our purpose (Table S6).

A

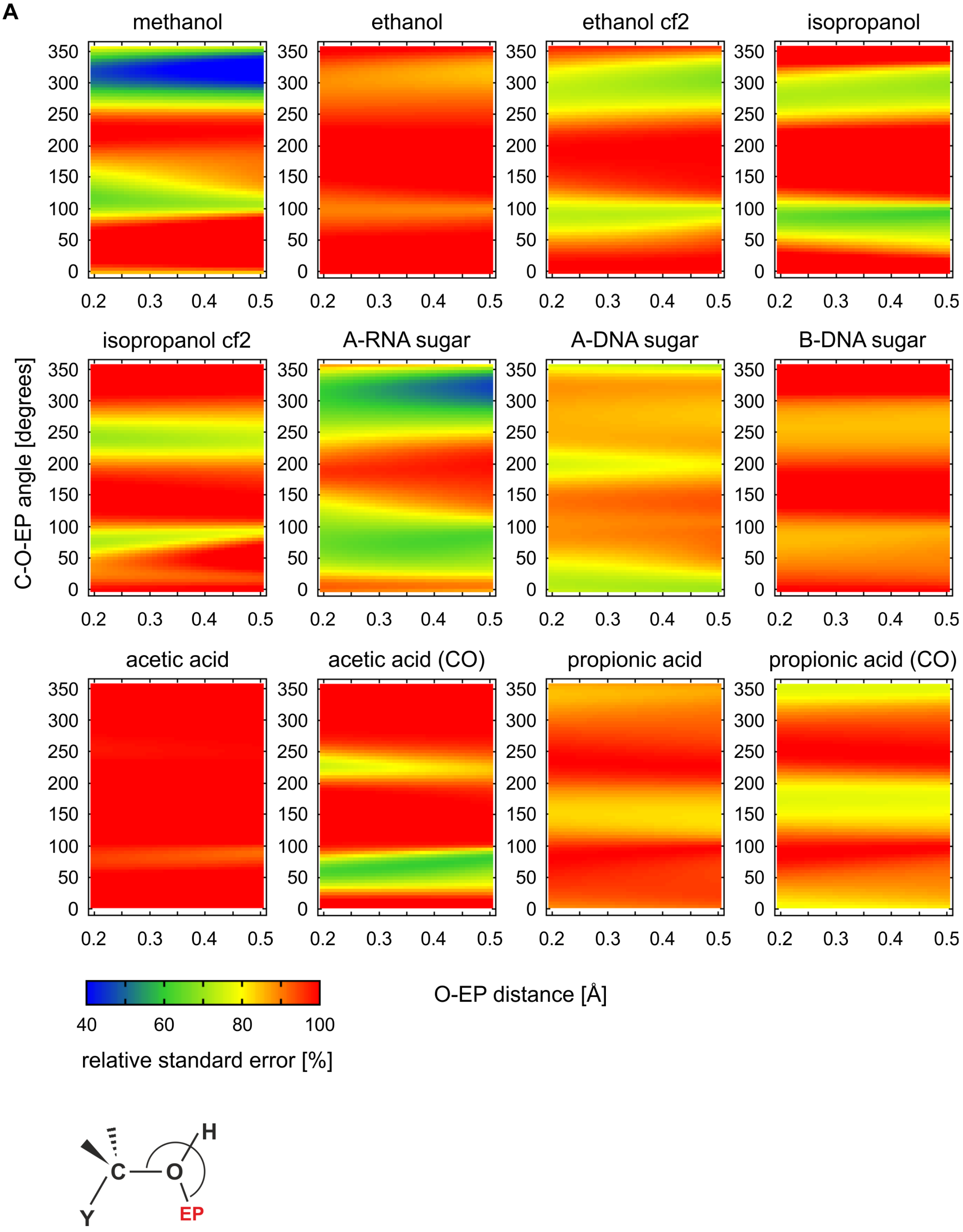
methanol
ethanol
ethanol cf2
isopropanol
isopropanol cf2
A-RNA sugar
A-DNA sugar
B-DNA sugar
acetic acid
acetic acid (CO)
propionic acid
propionic acid (CO)
C-O-EP angle [degrees]
0
50
100
150
200
250
300
350
0.2
0.3
0.4
0.5
O-EP distance [Å]
40
60
80
100
relative standard error [%]
H
C
O
Y
EP

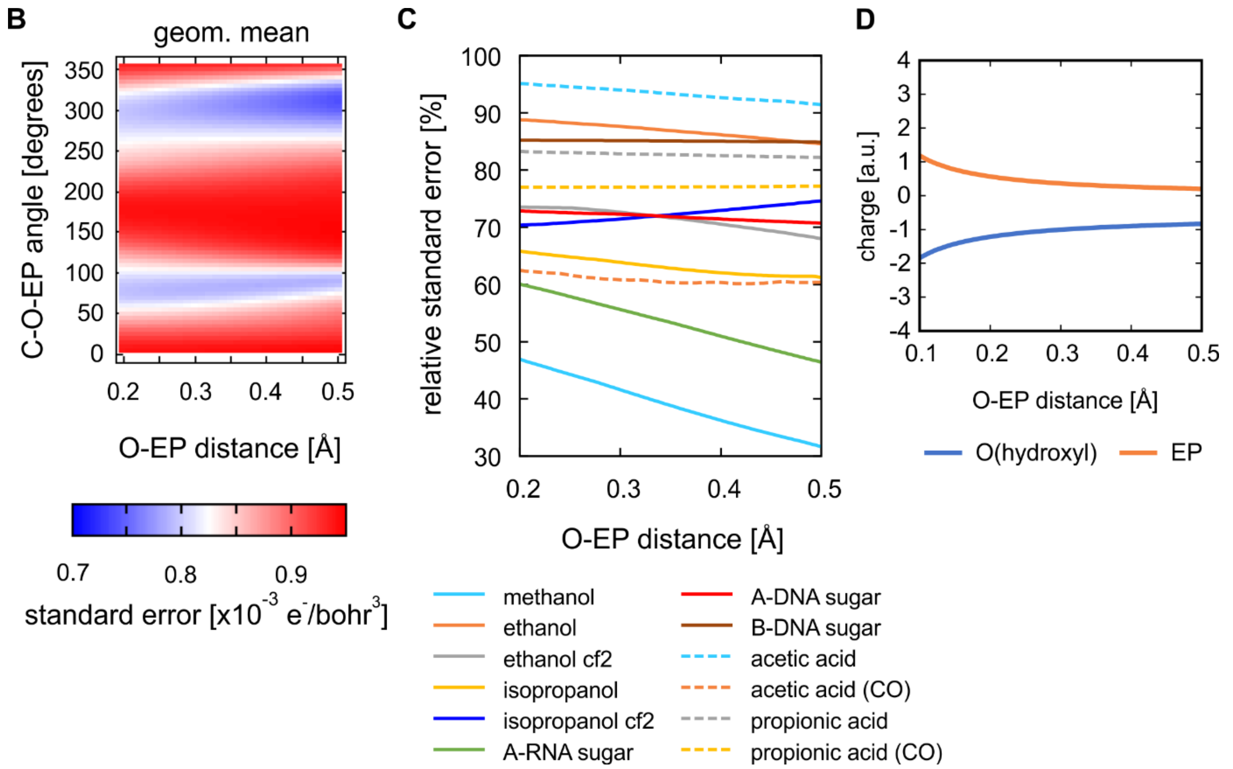

**Figure S3 – A**) Impact of position of 1 EP located on the hydroxyl group (O-EP distance and C-O-EP angle in the C-O-H plane) on the relative standard error (relative to the one without EPs) of ESP fit equivalenced in 2 stages of all compounds with hydroxyl group from the training set shown on 2D maps **B)** geometric mean of the standard error [x$10^{-3}$ Hartree/$e^{-}$] of all the compounds **C)** How relative standard error depends on O-EP distance (C-O-EP angle is always optimal for the given distance) **D)** Charge magnitudes of hydroxyl oxygen and EP of methanol as they depend on O-EP distance at the C-O-EP angle of 290°.

**Table S5** – Impact of including 1 EP on oxygen atom of hydroxyl group (O-EP distance=0.35Å, C-O-EP angle=290°, EP lies at the C-O-H plane, see Figure 3 in the Main text) on standard error of the ESP fit equivalenced in 2 stages and on minimal and maximal $V_i^{MM}$-$V_i^{QM}$ (all: ×$10^{-3}$ Hartree/$e^{-}$).

| | no EPs on hydroxyl | | | 1 EP per hydroxyl (ours, angle=290°) | | | Relative Std Error (EP:no EP) [%] |
|---|---|---|---|---|---|---|---|
| | Std Error | Min $V_i^{MM}$-$V_i^{QM}$ | Max $V_i^{MM}$-$V_i^{QM}$ | Std Error | Min $V_i^{MM}$-$V_i^{QM}$ | Max $V_i^{MM}$-$V_i^{QM}$ | |
| **methanol** | 1.756 | -10.664 | 5.623 | 0.961 | -6.064 | 3.444 | 54.7 |
| **ethanol** | 0.845 | -4.609 | 5.567 | 0.750 | -6.167 | 4.083 | 88.7 |
| **ethanol cf2** | 1.608 | -9.228 | 7.403 | 1.157 | -7.930 | 3.945 | 72.0 |
| **isopropanol** | 1.154 | -7.813 | 6.084 | 0.832 | -8.331 | 4.298 | 72.1 |
| **isopropanol cf2** | 0.565 | -5.433 | 3.119 | 0.518 | -6.498 | 2.394 | 91.6 |
| **A-RNA sugar** | 1.834 | -11.458 | 10.826 | 1.094 | -6.980 | 5.491 | 59.6 |
| **A-DNA sugar** | 1.015 | -7.530 | 6.957 | 0.870 | -7.063 | 5.648 | 85.7 |
| **B-DNA sugar** | 1.141 | -7.780 | 5.597 | 1.010 | -7.947 | 4.453 | 88.5 |
| **acetic acid** | 0.808 | -5.329 | 6.572 | 0.820 | -5.298 | 6.544 | 101.4 |
| **acetic acid (CO)** | 0.491 | -4.534 | 2.061 | 0.494 | -5.171 | 1.886 | 100.6 |
| **propionic acid** | 0.715 | -5.538 | 5.791 | 0.664 | -5.444 | 5.691 | 92.8 |
| **propionic acid (CO)** | 0.529 | -5.285 | 3.330 | 0.504 | -3.981 | 3.593 | 95.2 |

**(CO)=including 2 EPs on carbonyl group (see Figure 3 in the Main text)**
**cf2=conformer 2 (second most stable conformer)**

**Table S6 –** Impact of including 2 EPs on oxygen atom of hydroxyl group (placed at sp3-like position suggested by Dixon et al.[1]) on standard error of the ESP fit equivalenced in 2 stages and on minimal and maximal $V_i^{MM}$-$V_i^{QM}$ (all: $\times10^{-3}$ Hartree/e$^-$).

| | 2 EPs per hydroxyl (Dixon, sp3-like) | | | Relative Std Error (EP:no EP) [%] |
|---|---|---|---|---|
| | Std Error | Min $V_i^{MM}$-$V_i^{QM}$ | Max $V_i^{MM}$-$V_i^{QM}$ | |
| **methanol** | 1.830 | -10.924 | 6.819 | 104.2 |
| **ethanol** | 0.857 | -4.932 | 5.217 | 101.3 |
| **ethanol cf2** | 1.536 | -9.088 | 6.306 | 95.6 |
| **isopropanol** | 1.108 | -7.958 | 4.730 | 96.0 |
| **isopropanol cf2** | 0.500 | -5.936 | 3.130 | 88.4 |
| **A-RNA sugar** | 1.631 | -10.941 | 7.188 | 89.0 |
| **A-DNA sugar** | 0.924 | -7.570 | 6.150 | 91.0 |
| **B-DNA sugar** | 1.044 | -7.837 | 5.866 | 91.5 |
| **acetic acid** | 0.825 | -5.682 | 6.460 | 102.1 |
| **acetic acid (CO)** | 0.846 | -5.796 | 5.988 | 102.8 |
| **propionic acid** | 0.715 | -5.540 | 5.810 | 100.0 |
| **propionic acid (CO)** | 0.698 | -6.024 | 5.099 | 100.1 |

**(CO)=including 2 EPs on carbonyl group (at position suggested by Dixon et al.[1])**
**cf2=conformer 2 (second most stable conformer)**

*Thiol*

Optimal number and position of EPs on sulfur in organic compounds was thoroughly explored by Yan et al.[2] which recommend 2 symmetric EPs with S-EP distance of 0.7Å, C-S-EP angle of 96.8° and H-C-S-EP dihedral angle of 97.4° (i.e. H-S-EP angle of 96.46° and EP-S-EP angle of 160°) as consensus for all sulfur organic compounds. Similar position was also suggested previously by Dixon et al.[1]

Based on our data, we note that one EP only (located at S-EP distance of 0.7Å, C-S-EP angle of 280° at C-S-H plane) is sufficient to significantly improve the description of ESP around a thiol group (Table S7, Figures S4 A,B). However, the two symmetric EPs on the S atom suggested by Dixon et al.[1] and Yan et al.[2] provide greater reduction of standard error (Table S8, Figure S5A).

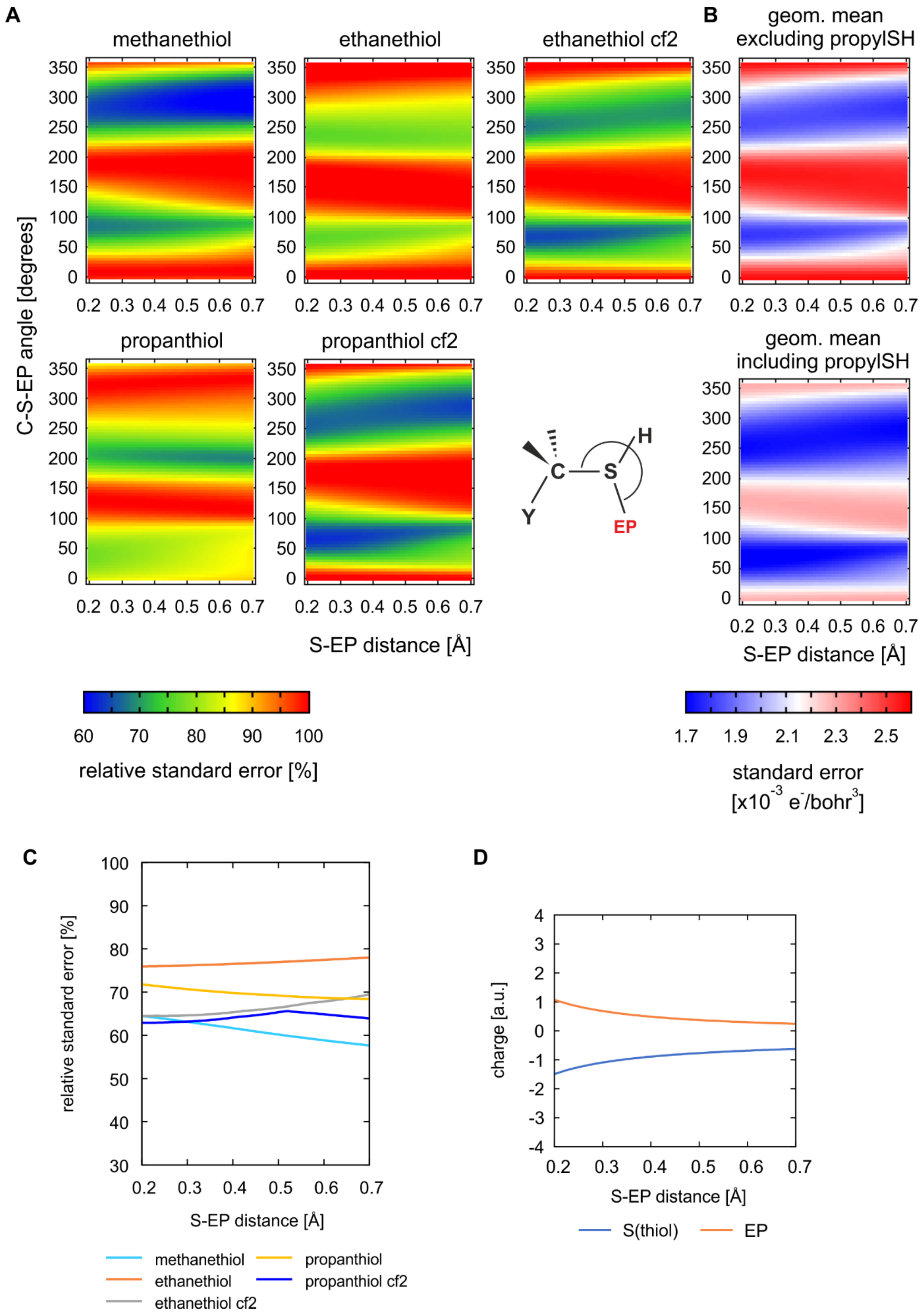

**Figure S4 - A**) Impact of position of 1 EP located on the thiol group (S-EP distance and C-S-EP angle in the C-S-H plane) on the relative standard error (relative to the one without EPs) of ESP fit equivalenced in 2 stages of all compounds with thiol group from the training set shown on 2D maps **B)** geometric mean of the standard error [x$10^{-3}$ Hartree/$e^{-}$] of all the compounds **C)** How relative standard error depends on S-EP distance (C-S-EP angle is always optimal for the given distance) **D**) Charge magnitudes of thiol sulfur and EP of methanethiol as they depend on S-EP distance at given C-S-EP angle of 280°.

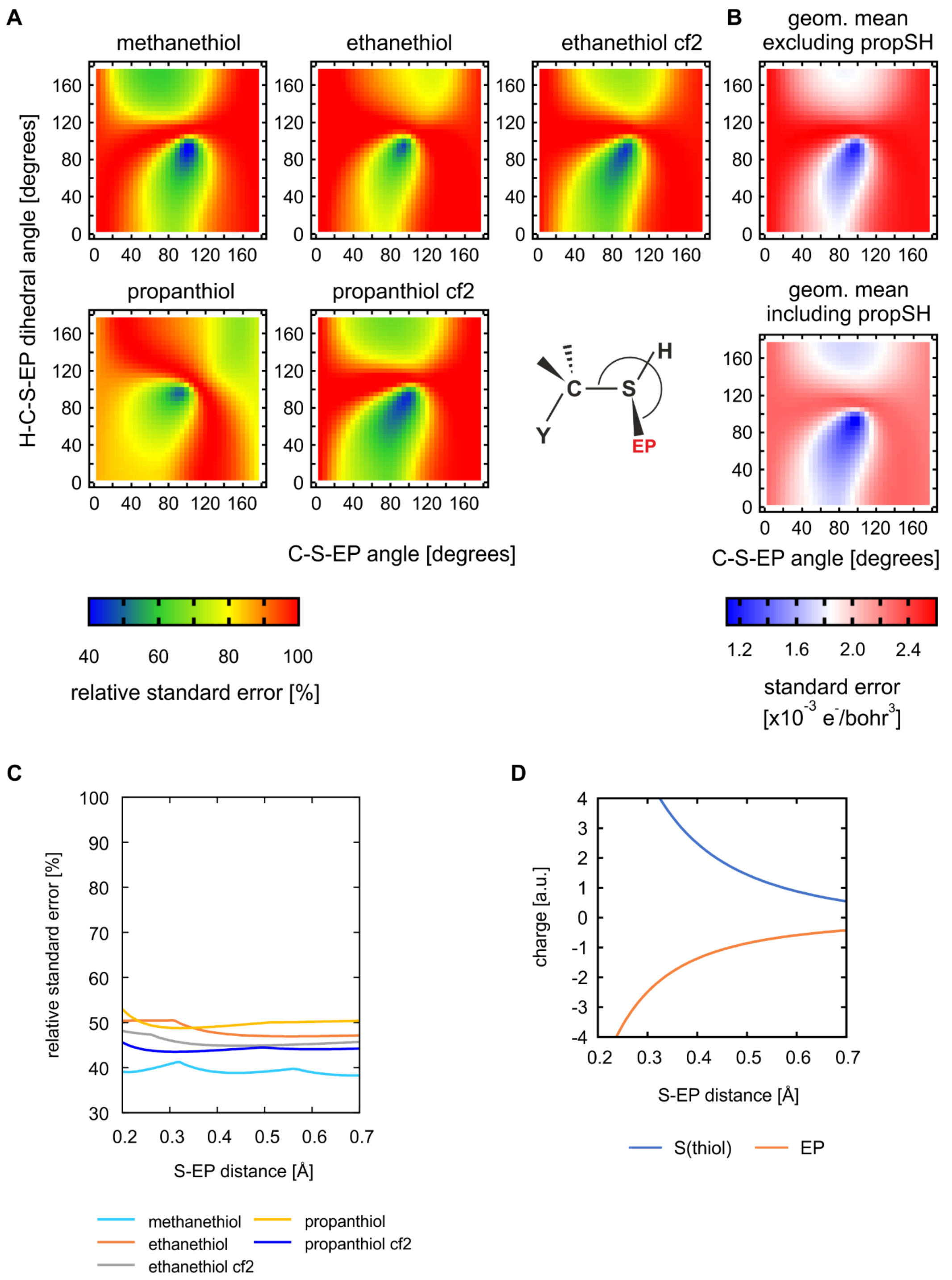

**Figure S5 - A**) Impact of position of 2 symmetric EPs located on the thiol group (C-S-EP angle and the H-C-S-EP dihedral angle whereas S-EP distance is 0.7Å) on the relative standard error (relative to the one without EPs) of ESP fit equivalenced in 2 stages of all compounds with thiol group from the training set shown on 2D maps **B)** geometric mean of the standard error [$x10^{-3}$ Hartree/$e^{-}$] of all the compounds **C)** How relative standard error depends on S-EP distance (C-S-EP angle and H-C-S-EP dihedral are always optimal for the given distance) **D**) Charge magnitudes of thiol sulfur and EP of methanethiol as they depend on S-EP distance at given C-S-EP angle of 95° and H-C-S-EP dihedral angle of 90°.

**Table S7** – Impact of including 1 EP on sulfur atom of thiol group (S-EP distance=0.70Å, C-S-EP angle=280° at C-S-H plane) on standard error of the ESP fit equivalenced in 2 stages and on minimal and maximal $V_i^{MM}$-$V_i^{QM}$ (all: ×10$^{-3}$ Hartree/e$^{-}$).

| | no EPs | | | 1 EP per thiol (ours) | | | Relative Std Error (EP:no EP) [%] |
|---|---|---|---|---|---|---|---|
| | Std Error | Min $V_i^{MM}$-$V_i^{QM}$ | Max $V_i^{MM}$-$V_i^{QM}$ | Std Error | Min $V_i^{MM}$-$V_i^{QM}$ | Max $V_i^{MM}$-$V_i^{QM}$ | |
| **methanethiol** | 3.025 | -25.259 | 12.801 | 1.786 | -16.634 | 7.889 | 59.0 |
| **ethanethiol** | 1.989 | -15.194 | 12.334 | 1.661 | -16.331 | 9.014 | 83.5 |
| **ethanethiol cf2** | 2.772 | -23.035 | 14.496 | 1.927 | -17.502 | 9.028 | 69.5 |
| **propanethiol** | 1.612 | -12.848 | 10.192 | 1.469 | -14.903 | 8.484 | 91.1 |
| **propanethiol cf2** | 2.629 | -23.570 | 13.440 | 1.687 | -16.789 | 7.748 | 64.2 |

**Table S8** – Impact of including 2 EPs on sulfur atom of thiol group as suggested by Dixon et al.[1] (S-EP distance=0.70Å, C-S-EP angle=90°, H-C-S-EP dihedral angle=90°) and Yan et al.[2] (S-EP distance=0.70Å, C-S-EP angle=96.8°, H-C-S-EP dihedral angle=97.4°) on standard error of the ESP fit equivalenced in 2 stages and on minimal and maximal $V_i^{MM}$-$V_i^{QM}$ (all: ×10$^{-3}$ Hartree/e$^{-}$).

| | 2 EPs per thiol (Dixon) | | | Relative Std Error (EP:no EP) [%] | 2 EPs per thiol (Yan) | | | Relative Std Error (EP:no EP) [%] |
|---|---|---|---|---|---|---|---|---|
| | Std Error | Min $V_i^{MM}$-$V_i^{QM}$ | Max $V_i^{MM}$-$V_i^{QM}$ | | Std Error | Min $V_i^{MM}$-$V_i^{QM}$ | Max $V_i^{MM}$-$V_i^{QM}$ | |
| **methanethiol** | 1.694 | -13.178 | 5.096 | 56.0 | 1.547 | -10.390 | 5.286 | 51.1 |
| **ethanethiol** | 1.014 | -8.193 | 3.647 | 51.0 | 0.941 | -6.310 | 4.822 | 47.3 |
| **ethanethiol cf2** | 1.357 | -10.297 | 4.987 | 49.0 | 1.365 | -8.260 | 8.036 | 49.3 |
| **propanethiol** | 0.871 | -8.983 | 4.355 | 54.0 | 0.808 | -6.994 | 4.598 | 50.1 |
| **propanethiol cf2** | 1.309 | -10.343 | 6.441 | 49.8 | 1.341 | -8.333 | 8.509 | 51.0 |

*Amine*

One EP at counter-intuitive position (with C-N-EP angle of 25°) can significantly improve the standard error of the ESP fit (Figure S6A) but it leads to EP and N charges unsuitable for MD (Figure S6D). Thus, if one wants to use 1 EP we would rather recommend location with C-N-EP angle of 225° (Figure S6D). However, the best improvement is achieved with two EPs (Figure S7A, Table S9). The position that provides the largest reduction of standard error (C-N-EP2 angle=290°) leads to absolute charges values reaching 2 a.u. at the N-EP distance of 0.35Å (Figure S7D). This issue can be solved by moving EP2 slightly.

One EP located near intuitive position of electron lone pair[1] improves only poorly the ESP fit (Table S10). As a check, the EP parameters were tested also on propylamine.

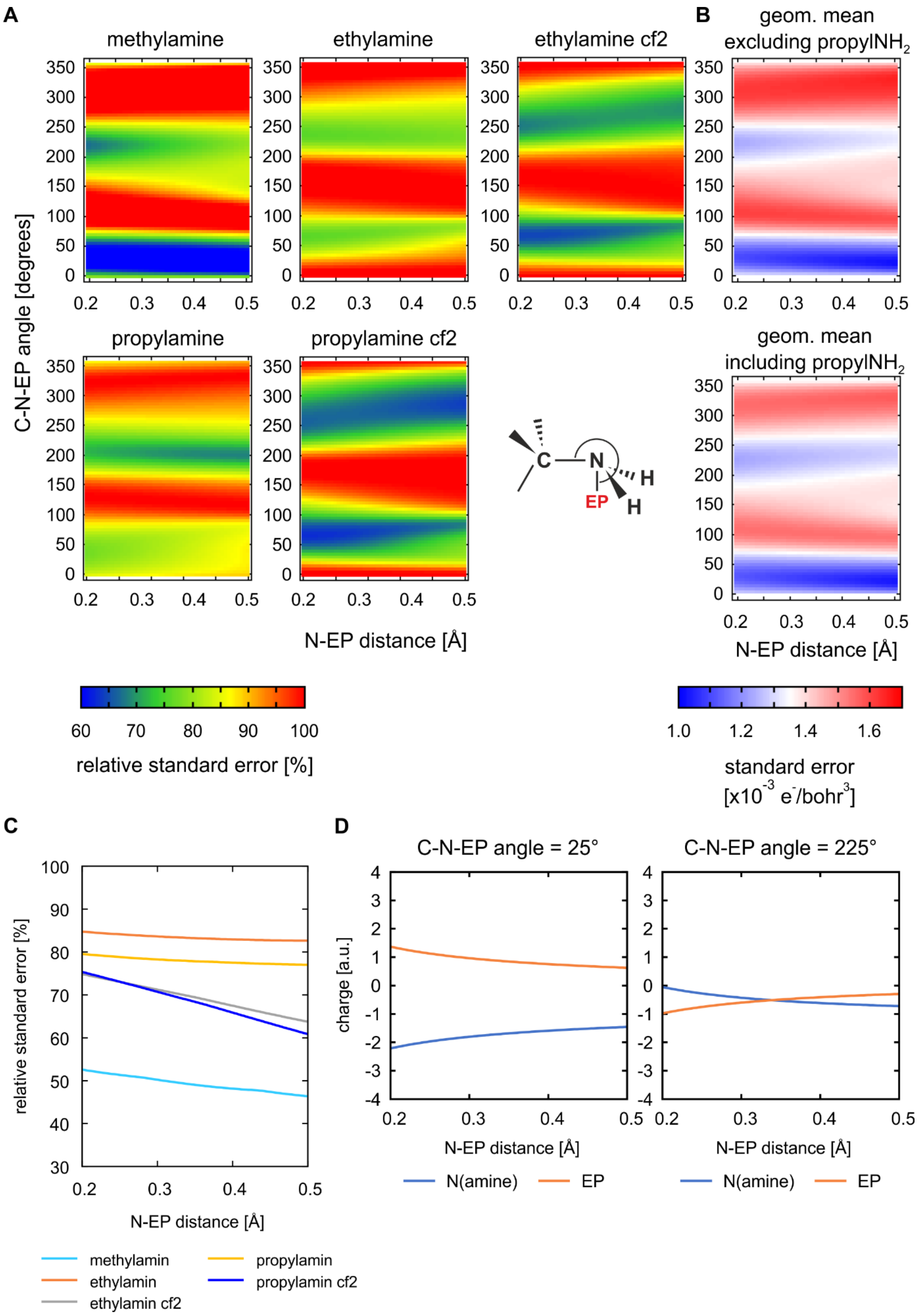

**Figure S6 - A**) Impact of position of 1 EP located on the amine group (N-EP distance and C-N-EP angle whereas the EP lies in the plane which bisects the amine group) on the relative standard error (relative to the one without EPs) of ESP fit equivalenced in 2 stages of all compounds with amine group from the training set shown on 2D maps **B)** geometric mean of the standard error [$x10^{-3}$ Hartree/$e^{-}$] of all the compounds **C)** How relative standard error depends on N-EP distance (C-N-EP angle is always optimal for the given distance) **D**) Charge magnitudes of amine nitrogen and EP of methylamine as they depend on N-EP distance for two given C-N-EP angles.

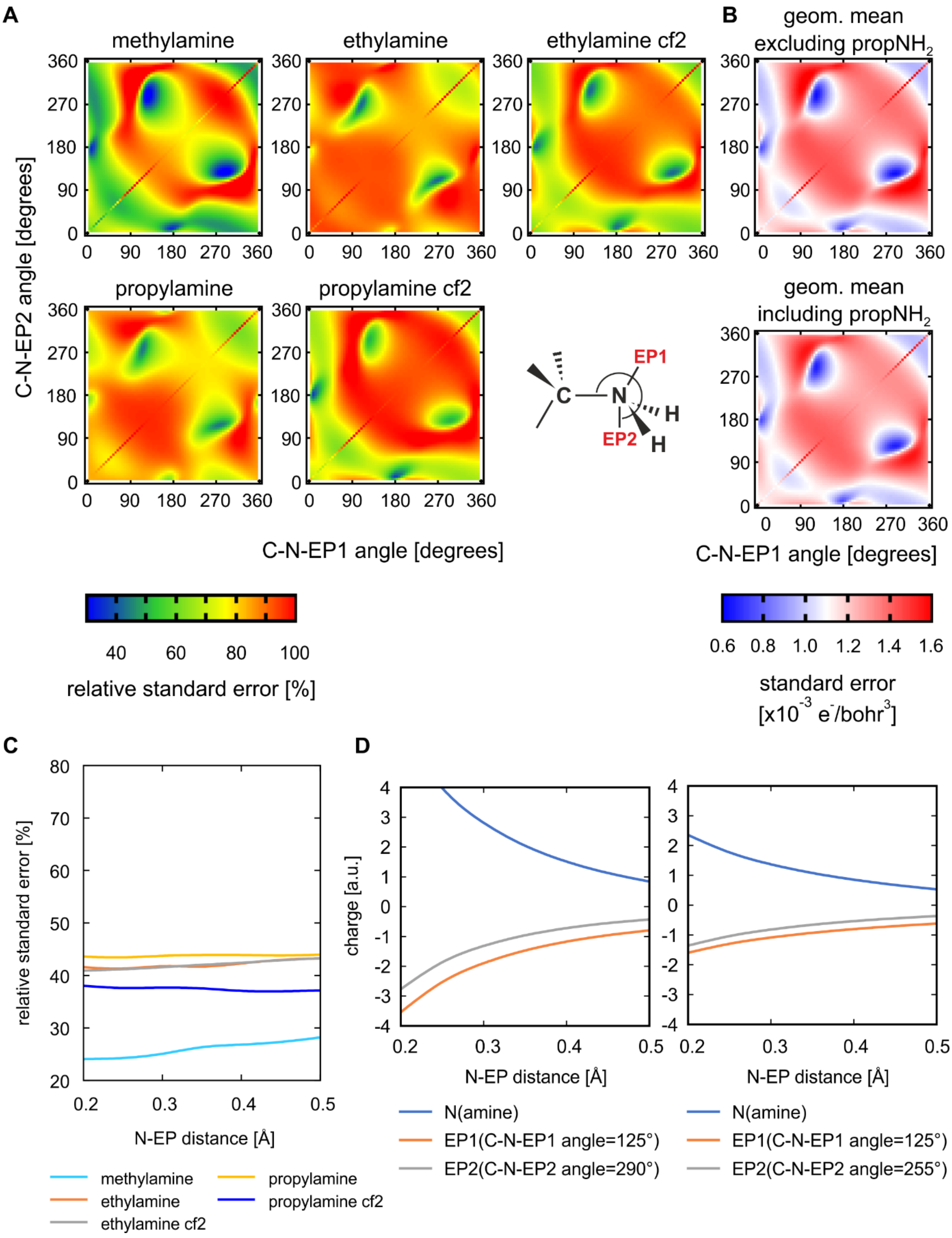

**Figure S7 - A**) Impact of position of 2 EPs located on the amine group (C-N-EP1 and EP2 angles at given N-EP distance of 0.35Å) on the relative standard error (relative to the one without EPs) of ESP fit equivalenced in 2 stages of all compounds with amine group from the training set shown on 2D maps **B)** geometric mean of the standard error [x$10^{-3}$ Hartree/$e^-$] of all the compounds **C)** How relative standard error depends on N-EP distance (C-N-EP1 and EP2 angles are always optimal for the given distance) **D**) Charge magnitudes of amine nitrogen and 2 EPs of methylamine as they depend on N-EP distance for the given C-N-EP1 angle and two different C-N-EP2 angles.

**Table S9** – Impact of including 2 EPs on nitrogen atom of amine group (N-EP distance=0.35Å, C-N-EP1 and C-N-EP2 angles = 125° and 255°, respectively, see Figure 3 in the Main text) on standard error of the ESP fit equivalenced in 2 stages and on minimal and maximal $V_i^{MM}$-$V_i^{QM}$ (all: $\times 10^{-3}$ Hartree/$e^-$).

| | no EPs | | | 2 EPs (ours) | | | Relative Std Error (EP:no EP) [%] |
|---|---|---|---|---|---|---|---|
| | **Std Error** | **Min $V_i^{MM}$-$V_i^{QM}$** | **Max $V_i^{MM}$-$V_i^{QM}$** | **Std Error** | **Min $V_i^{MM}$-$V_i^{QM}$** | **Max $V_i^{MM}$-$V_i^{QM}$** | |
| **methylamine** | 2.026 | -9.770 | 6.276 | 0.535 | -3.019 | 4.059 | 26.4 |
| **ethylamine** | 1.174 | -7.187 | 6.641 | 0.663 | -3.426 | 3.701 | 56.5 |
| **ethylamine cf2** | 1.576 | -9.726 | 6.161 | 0.689 | -4.028 | 3.657 | 43.7 |
| **propylamine** | 1.254 | -7.123 | 9.417 | 0.600 | -3.207 | 4.627 | 47.9 |
| **propylamine cf2** | 1.633 | -12.094 | 6.272 | 0.841 | -3.364 | 4.963 | 51.5 |

**Table S10** – Impact of including 1 EP on nitrogen atom of amine group (N-EP distance=0.35Å, C-N-EP angle = 109.5° as suggested by Dixon et al.[1]) on standard error of the ESP fit equivalenced in 2 stages and on minimal and maximal $V_i^{MM}$-$V_i^{QM}$ (all: $\times 10^{-3}$ Hartree/$e^-$).

| | 1 EP (sp3-like, Dixon) | | | Relative Std Error (EP:no EP) [%] |
|---|---|---|---|---|
| | **Std Error** | **Min $V_i^{MM}$-$V_i^{QM}$** | **Max $V_i^{MM}$-$V_i^{QM}$** | |
| **methylamine** | 2.275 | -8.019 | 12.899 | 112.3 |
| **ethylamine** | 1.099 | -5.244 | 9.578 | 93.7 |
| **ethylamine cf2** | 1.610 | -9.425 | 7.343 | 102.1 |
| **propylamine** | 1.243 | -6.321 | 10.833 | 99.1 |
| **propylamine cf2** | 1.708 | -11.590 | 7.443 | 104.6 |

*Ammonium*

One EP diminishes sufficiently standard error of both ammonium compounds from our training set, in particular under $7 \times 10^{-4}$ Hartree/$e^-$ (Figure S8B, Table S11). As a check, the EP parameters were tested also on propylammonium.

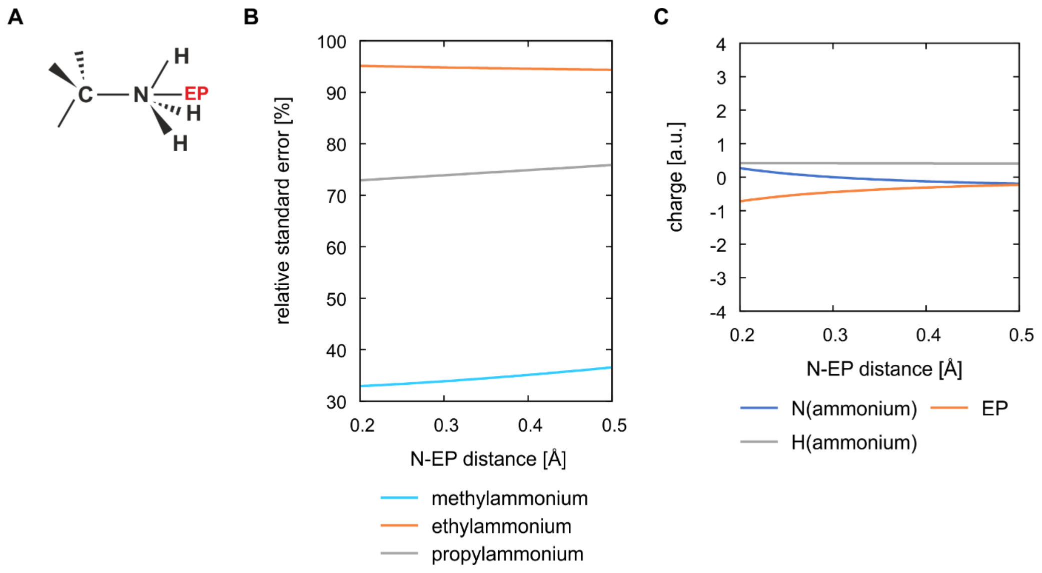

**Figure S8 – A)** Extra point is located on C-N axis. **B)** Relative standard error (with respect to model without EPs) of the ESP fit equivalenced in 2 stages as it depends on N-EP distance, of all appropriate compounds from our training set. **C)** Charge magnitudes of ammonium nitrogen, hydrogen and EP of methylammonium ion as they depend on N-EP distance.

**Table S11** – Impact of including 1 EP on nitrogen atom of ammonium group (on the C-N axis with N-EP distance of 0.35Å) on standard error of the ESP fit equivalenced in 2 stages, and on minimal and maximal $V_i^{MM}$-$V_i^{QM}$ (all: ×$10^{-3}$ Hartree/$e^{-}$).

| | no EPs | | | 1 EP | | | Relative Std Error (EP:no EP) [%] |
|---|---|---|---|---|---|---|---|
| | **Std Error** | **Min $V_i^{MM}$-$V_i^{QM}$** | **Max $V_i^{MM}$-$V_i^{QM}$** | **Std Error** | **Min $V_i^{MM}$-$V_i^{QM}$** | **Max $V_i^{MM}$-$V_i^{QM}$** | |
| **methylammonium** | 0.852 | -3.243 | 4.685 | 0.294 | -2.338 | 1.694 | 34.5 |
| **ethylammonium** | 0.678 | -2.826 | 5.391 | 0.642 | -3.066 | 4.738 | 94.7 |
| **propylammonium** | 0.925 | -3.342 | 5.930 | 0.688 | -2.762 | 5.948 | 74.4 |

*Bridging oxygen*

The only compound from the training set carrying bridging oxygen is dimethylether. We examined position of two symmetric EPs located either in the C-O-C plane or in the plane perpendicular to the C-O-C plane and crossing bisector of C-O-C angle (Figures 9A,E). When located in the C-O-C plane, the 2 EPs improves standard error of the ESP fit very significantly (Figure S9F) but also lead to very high absolute values of charges (Figure S9H). Therefore, the only usable position of two symmetric EPs is in the plane perpendicular to the C-O-C plane where they only moderately diminish the standard error (Figure S9B, Table S12) but keep reasonable values of charges (Figure S9D).

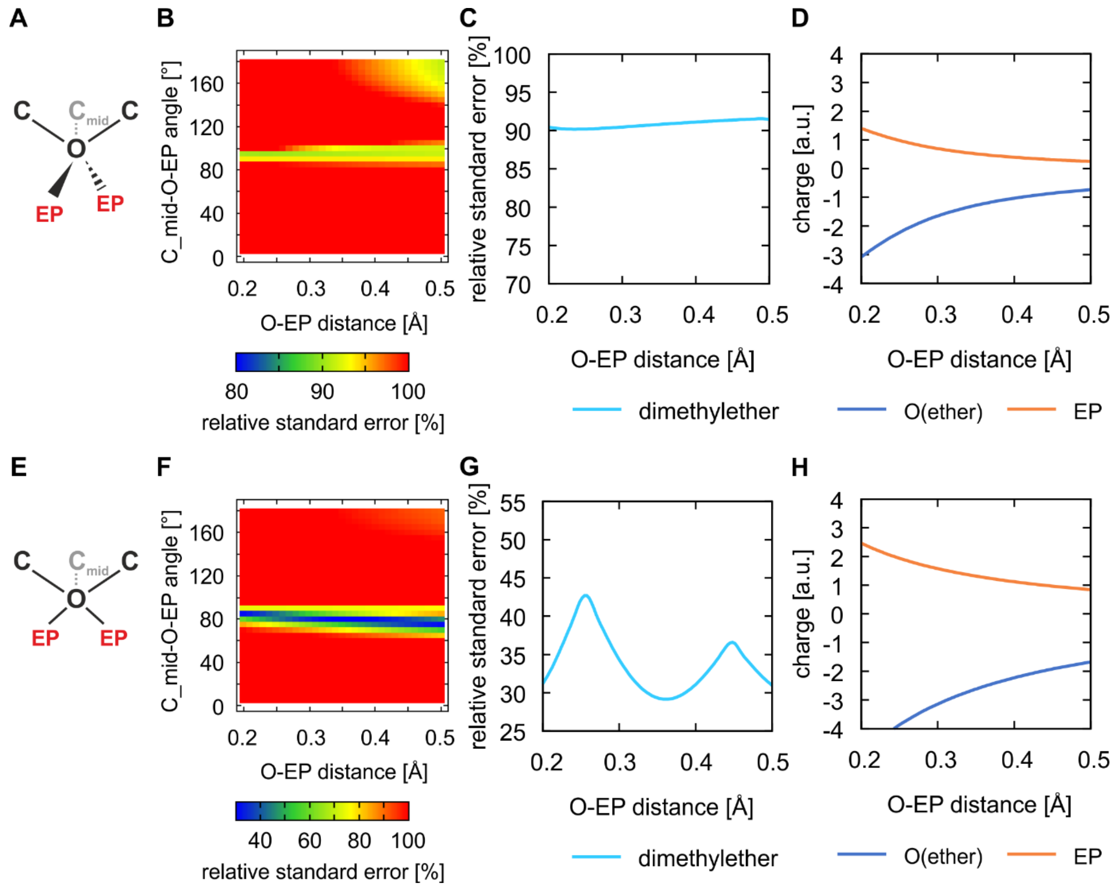

**Figure S9 – A)** Two symmetric EPs are located in the plane which is perpendicular to the C-O-C plane and crosses C-O-C bisector **B**) Impact of position (O-EP distance and $C_{mid}$-O-EP angle) of the 2 EPs located in this plane on the relative standard error (normalized to model without EPs) of ESP fit equivalenced in 2 stages, of dimethylether shown on 2D map **C)** How relative standard error depends on O-EP distance ($C_{mid}$-O-EP angle is always optimal for the given distance) **D**) Charge magnitudes of ether oxygen and EP of dimethylether as they depend on O-EP distance for the $C_{mid}$-O-EP angle of 95°. **E), F), G), H)** 2 symmetric EPs are located in the C-O-C plane

**Table S12** – Impact of including ours EPs to ether, thioether and secondary amine groups on standard error of the ESP fit equivalenced in 2 stages and on minimal and maximal $V_i^{MM}$-$V_i^{QM}$ (all: ×$10^{-3}$ Hartree/$e^-$).

| | no EPs | | | ours EPs | | | Relative Std Error (EP:no EP) [%] |
|---|---|---|---|---|---|---|---|
| | **Std Error** | **Min $V_i^{MM}$-$V_i^{QM}$** | **Max $V_i^{MM}$-$V_i^{QM}$** | **Std Error** | **Min $V_i^{MM}$-$V_i^{QM}$** | **Max $V_i^{MM}$-$V_i^{QM}$** | |
| **dimethylether** | 1.057 | -4.119 | 7.616 | 0.960 | -4.219 | 9.589 | 90.8 |
| **dimethylthioether** | 1.749 | -14.586 | 12.056 | 1.162 | -6.626 | 5.193 | 66.4 |
| **dimethylamine** | 1.971 | -14.429 | 5.015 | 1.275 | -7.933 | 5.326 | 64.7 |

**Table S13** – Impact of including EPs suggested by Dixon et al.[1] to ether, thioether and secondary amine groups on standard error of the ESP fit equivalenced in 2 stages and on minimal and maximal $V_i^{MM}$-$V_i^{QM}$ (all: ×$10^{-3}$ Hartree/$e^-$).

| | EPs (Dixon) | | | Relative Std Error (EP:no EP) [%] |
|---|---|---|---|---|
| | **Std Error** | **Min $V_i^{MM}$-$V_i^{QM}$** | **Max $V_i^{MM}$-$V_i^{QM}$** | |
| **dimethylether** | 1.331 | -4.848 | 6.114 | 125.9 |
| **dimethylthioether** | 1.377 | -8.556 | 6.292 | 78.7 |
| **dimethylamine** | 2.634 | -6.002 | 11.976 | 133.7 |

*Bridging sulfur*

The only compound from the training set carrying bridging sulfur is dimethylthioether. We examined position of two symmetric EPs located in the plane perpendicular to the C-S-C plane (Figure S10A, Table S12). The optimal position of the EPs lies at $C_{mid}$-S-EP angle of 110° but this also leads to absolute values of charges greater than 1 a.u. Acceptable values of charges can be obtained when EPs are moved to $C_{mid}$ -S-EP angle of 100° at distance of 0.7Å. This position fully agrees with the consensus identified by Yan et al.[2]

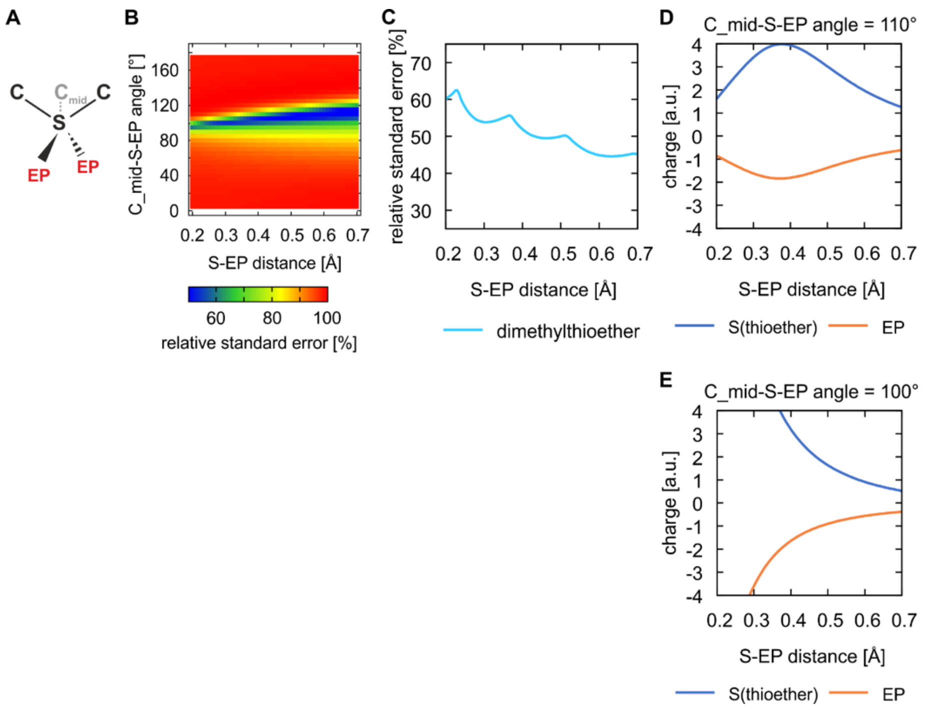

**Figure S10 – A)** Two symmetric EPs of the thioether group are located in the plane which is perpendicular to the C-S-C plane and crosses the bisector of C-S-C angle **B**) Impact of S-EP distance and $C_{mid}$-S-EP angle of the two symmetric EPs on the relative standard error (normalized to the model without EPs) of ESP fit equivalenced in 2 stages, of dimethylthioether shown on 2D map **C)** How relative standard error depends on S-EP distance ($C_{mid}$-S-EP angle is always optimal for the given distance) **D**) Charge magnitudes of thioether sulfur and EP of dimethylthioether as they depend on S-EP distance for the $C_{mid}$-S-EP angle of 110° and **E)** 100°.

*Secondary amine*

The only compound from the training set carrying secondary amine group is dimethylamine. The optimal position of one EP for the compound lies in the plane which separates the amine group to two symmetric halves and has H-N-EP angle of 190° (Figures S11A,B, Table S12).

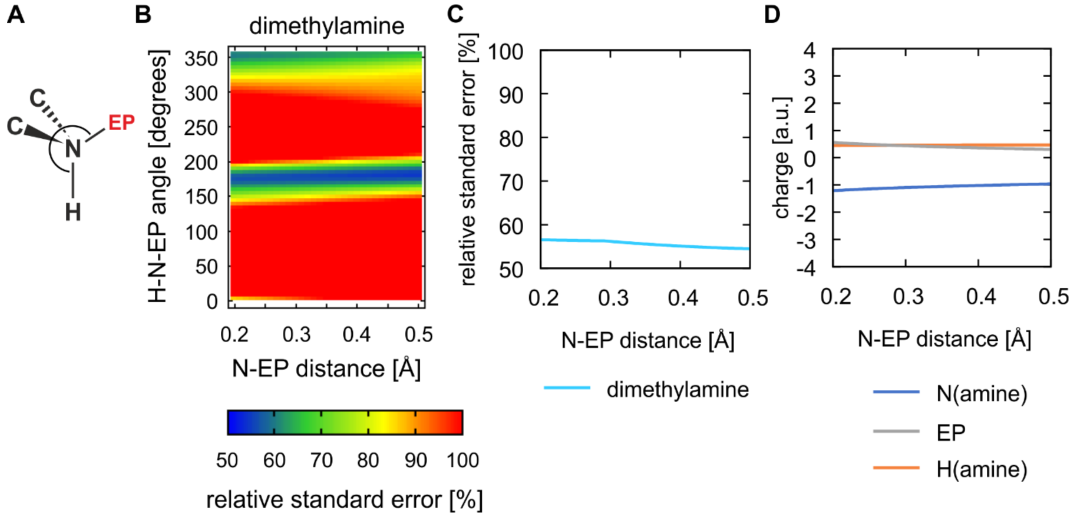

**Figure S11 – A)** One EP of the secondary amine group is located in the plane perpendicular to the C-N-C plane and crossing the bisector of the C-N-C angle. **B)** Impact of the N-EP distance and the H-N-EP angle on the relative standard error (normalized to the model without EPs) of ESP fit equivalenced in 2 stages, of dimethylamine shown on 2D map **C)** How relative standard error depends on N-EP distance (H-N-EP angle is always optimal for the given distance) **D)** Charge magnitudes of amine nitrogen, hydrogen and EP of dimethylamine as they depend on N-EP distance for the H-N-EP angle of 190°.

*Phosphate*

The ESP around the phosphate ion or its methyl derivatives is too complex that the ESP fit cannot be improved by adding few simple EPs.

**Optimization of the W-RESP scaling coefficient**

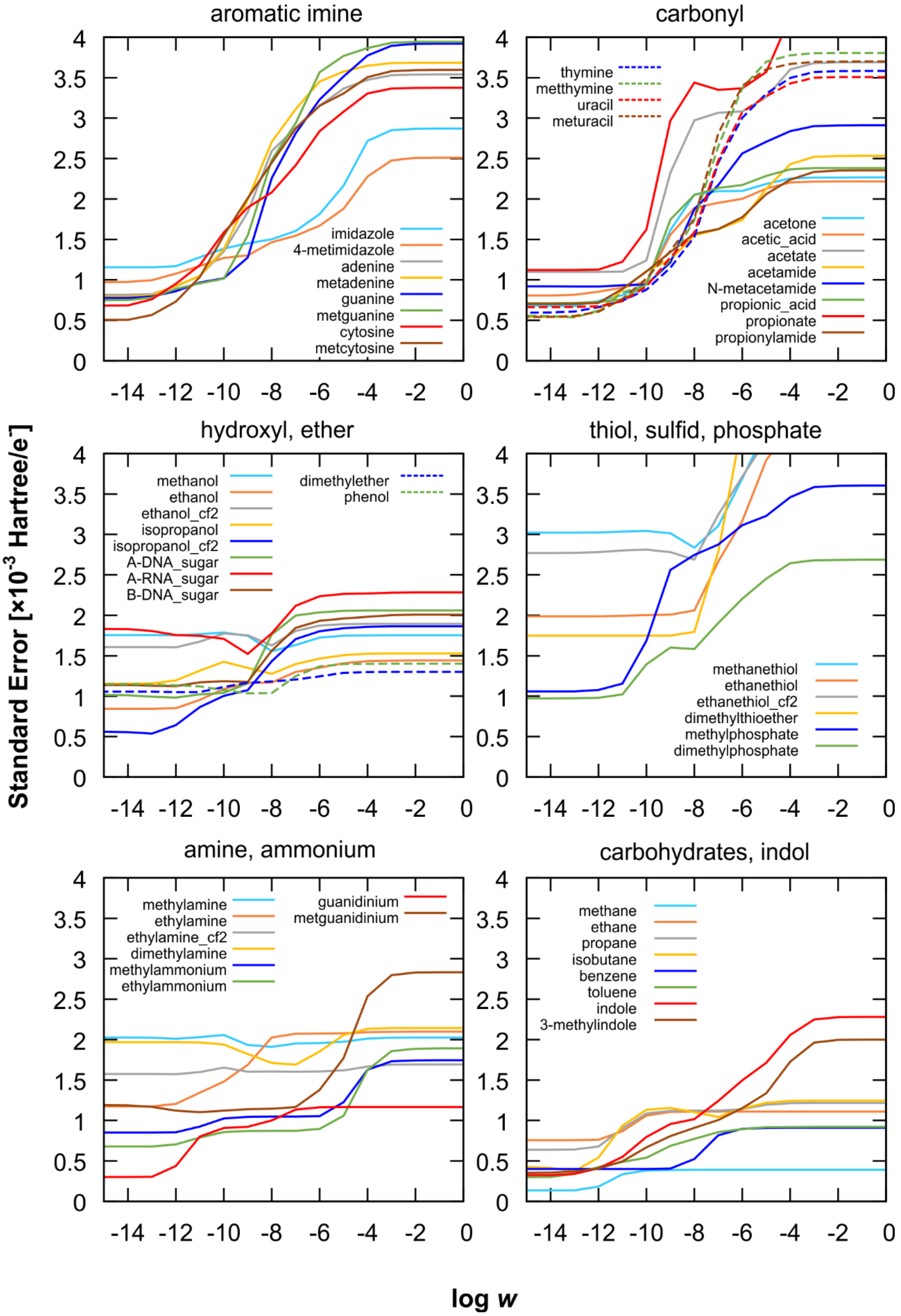

**Figure S12** – Impact of the W-RESP restraint with force constant ***w*** on the standard error of the ESP fit of the classical atom-centered charges. 47 compounds from our training set are grouped according to their functional groups.

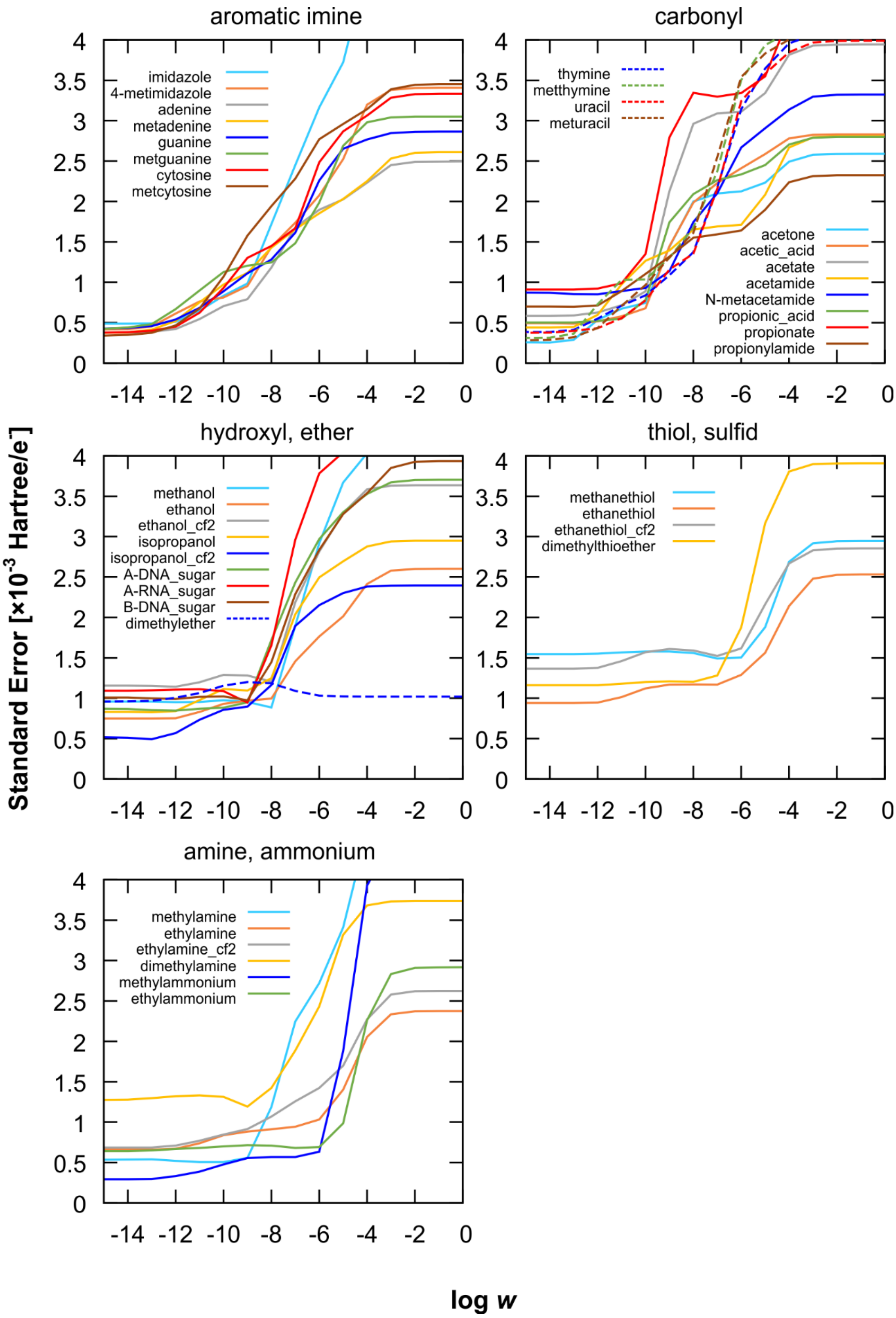

**Figure S13** – Impact of the W-RESP restraint with force constant ***w*** on the standard error of the ESP fit including extra points.

### Optimized glycosidic torsion parameters for W-RESP and W-RESP-EP charges

**Table S14** – Optimized χ dihedral terms for the revised W-RESP and W-RESP-EP charges, i.e., the O4'-C1'-N1-C8 and O4'-C1'-N1-C6 torsion angles for purine and pyrimidine nucleotides, respectively. The force constants ($k$) are in kcal·mol$^{-1}$ and the phase angles ($\theta$) are in degrees.

| | | | | W-RESP | | | | |
|---|---|---|---|---|---|---|---|---|
| Nucleotide | $k_1$ | $k_2$ | $k_3$ | $k_4$ | $\theta_1$ | $\theta_2$ | $\theta_3$ | $\theta_4$ |
| A | 1.07831 | 1.00558 | 0.42375 | 0.29764 | 105.4569 | 20.9530 | 172.0995 | 23.8573 |
| G | 1.09973 | 1.00826 | 0.37953 | 0.22295 | 105.2197 | 4.8602 | 161.2306 | 6.6159 |
| C | 0.96535 | 1.64450 | 0.90797 | 0.28870 | 138.0432 | 14.3599 | 189.9984 | 41.8949 |
| U/T | 1.35480 | 1.75082 | 0.58319 | 0.34269 | 151.0094 | 16.7928 | 172.8482 | 7.0541 |
| | | | | W-RESP-EP | | | | |
| Nucleotide | $k_1$ | $k_2$ | $k_3$ | $k_4$ | $\theta_1$ | $\theta_2$ | $\theta_3$ | $\theta_4$ |
| A | 1.13316 | 0.90769 | 0.51758 | 0.26257 | 125.2961 | 21.4722 | 165.5684 | 18.2682 |
| G | 1.14386 | 0.83758 | 0.45521 | 0.20098 | 124.4698 | 8.9705 | 159.1667 | 6.4998 |
| C | 1.02961 | 1.50054 | 0.83631 | 0.42015 | 113.8865 | 36.0486 | 185.8743 | 43.5772 |
| U/T | 1.26485 | 1.57903 | 0.49895 | 0.34533 | 138.8092 | 29.0303 | 169.3718 | 18.1136 |
